# Changes in the functional diversity and abundance of ectomycorrhizal fungi are decoupled from water uptake patterns in European beech forests

**DOI:** 10.1101/2024.07.09.602659

**Authors:** Asun Rodríguez-Uña, David Moreno-Mateos, Silvia Matesanz, Lisa Wingate, Adrià Barbeta, Javier Porras, Teresa E. Gimeno

## Abstract

Temperate forests on their warm and dry distribution limit are expected to be most vulnerable to reductions in water availability. This prediction is mostly based on studies assessing single forest functions, mainly growth. Water and nutrient cycling are functions that rely on tree roots and their symbiotic association with ectomycorrhizal (ECM) fungi. Trees can compensate for seasonal reductions in water availability by shifting root water-uptake (RWU) towards deeper soil layers, but ECM fungi dwell in the upper soil, thus suffering from desiccation and compromising nutrient uptake. We hypothesised that drier sites should depict larger seasonal shifts in RWU, but at the expense of lower diversity and colonization of fine roots by ECM fungi. We selected three beech (*Fagus sylvatica*) forests in their warm distribution limit with contrasting geographic locations and mean annual precipitation: northern Atlantic (2500mm), intermediate transitional (1150mm) and southern Mediterranean (780mm). We collected soil, stem and root samples in spring (wet) and summer (dry) to quantify fine-root density and colonization by ECM fungi, to infer RWU from isotopic composition of plant and soil water, and to characterize ECM fungal diversity through DNA-metabarcoding. High moisture in the upper soil benefited the ECM community, but higher diversity and fine-root colonization by ECM fungi in the upper soil did not imply larger contributions of this soil layer to RWU. The prevailing climate and local abiotic conditions determined how ECM communities structured, more than seasonal variability. Across sites, ECM communities differed in their functional diversity: ECM fungi with long hyphae, more vulnerable to water scarcity, dominated at the site with the highest water availability. Our results suggest that transient reductions in soil water availability might not compromise RWU but could be detrimental for maintaining ECM-mediated nutrient uptake in beech forests experiencing longer and more severe drought periods under current climate change.

## Introduction

Climate change is increasing the frequency and severity of drought episodes worldwide (Dai 2013). Drought risk is now extending to regions not considered water limited, including European temperate forests. In the past two decades, European forests have suffered multiple severe droughts, with detrimental impacts for forest health and productivity (e.g. Camarero et al. 2015, Ciais et al. 2005, Schuldt et al. 2020, Senf et al. 2018, Treydte et al. 2023). European beech (*Fagus sylvatica* L.) forests spread across Europe and constitute one iconic example of this drought-related decline (Geßler et al. 2007, Leuschner 2020). Numerous studies have demonstrated the negative impacts of drought on beech performance, mainly based on decreased radial growth, but also on canopy vigour or ecophysiological performance (Martinez del Castillo et al. 2022, Mazza et al. 2024, Serra-Maluquer et al. 2019, Zimmermann et al. 2015). Such drought sensitivity implies that under climate change the distribution range of beech could contract along its southern and low elevation limits. Evidence for such range contraction and lower performance along the drier and warmer limits has been found for various beech forests (Martinez del Castillo et al. 2022, Peñuelas and Boada 2003, Rozas et al. 2015). In contrast, similar studies have shown that marginal beech populations or those inhabiting their rear-edge are not necessarily more drought sensitive (Bert et al. 2022, Vilà-Cabrera and Jump 2019, Weber et al. 2013). Most of these studies based their predictions on a single aboveground forest function, radial growth. Yet, drought and climate change impact all forest functions, including not only those aboveground, but also belowground.

Ectomycorrhizae, symbiotic associations between tree roots and fungi, are crucial belowground interactions for ecosystem functioning (Tedersoo et al. 2020, Van der Heijden et al. 2015). In this association, the ectomycorrhizal (ECM) fungi (mainly from the phyla Basidiomycota and Ascomycota) form a hyphal mantle around the tree roots, enhancing nutrient and water uptake and providing pathogen resistance for their host in exchange for carbohydrates (Rasmann et al. 2017, Smith and Read 2008). ECM hyphae improve nutrient uptake by exudating enzymes, organic acids and other compounds that mobilize N and P locked in the soil organic matter or that help weather mineral surfaces (Finlay 2008). ECM fungi should also enhance water uptake by increasing the soil volume explored through the extension of their hyphal mantle. However, the relative importance of ECM fungi to plant water uptake appears to depend on certain functional traits, mainly the exploration type of the fungal partner (Johnson 2018, Prieto et al. 2016). The pattern and extent of the hyphal network (short contact vs. long exploration types) determines the soil exploration capacity (Agerer 2001), with crucial implications for the ECM fungal community structure and functioning (Koide, Fernandez and Malcolm 2014). At the same time, factors like climate, past land use, soil pH, nutrient levels and specially water availability strongly influence the structure and composition of the ECM fungal community (Correia et al. 2021, Erlandson et al. 2016, Querejeta, Egerton-Warburton and Allen 2009, Reis et al. 2018, Rodríguez-Uña et al. 2024, Van der Linde et al. 2018). The hyphae of ECM fungi are particularly sensitive to water availability because they dwell in the upper soil, lack regulatory mechanisms to control for water loss and suffer indirectly from the effects of drought on their host (Brunner et al. 2015, Gehring, Swaty and Deckert 2017).

At the onset of drought, plants activate mechanisms to preserve their hydraulic system (Brunner et al. 2015, Magnani, Grace and Borghetti 2002). For example, by taking advantage of fine roots as hydraulic fuses that decouple the plant from the drying soil (Jackson, Sperry and Dawson 2000). To activate this mechanism, lacunae (air gaps) form in the cortical cells of fine roots, triggering loss of conductivity (Cuneo et al. 2016), but this mechanism inevitably affects their associated ECM fungi (Gehring, Swaty and Deckert 2017). Drought also impairs photosynthesis and phloem transport, and hence reduces the amount and type of carbohydrates trees export to their fungal partners (Köhler et al. 2018, Ruehr et al. 2009, Shi et al. 2002). Overall, drought decreases the abundance, diversity, biomass and enzymatic activity of ECM fungi (Gehring, Swaty and Deckert 2017, Nickel et al. 2018), inducing shifts in community composition (Shi et al. 2002). For example, drought may drive a shift in the ECM fungal community towards a higher portion of ECM associations with fungal partners with a lower C-demand, i.e. contact- and short-exploration ECM fungal types (Castaño et al. 2018, Fernandez et al. 2017). In contrast, several studies have reported that ECM symbiosis enhances drought tolerance by increasing water absorbing surface area; hence, under drought, ECM networks formed by fungi with long emanating tissues (i.e. medium- and long-exploration ECM fungal types) should be favoured, although these are generally more C-demanding (Brunner et al. 2015, Köhler et al. 2018, Leuschner 2020, Nickel et al. 2018). It is thus unclear which ECM functional types would be most resistant and beneficial for tree survival and growth under low water availability, as well as the extent of the contribution of ECM fungi to root water uptake.

Inferring tree root water uptake is challenging. Concurrent analyses of the isotopic composition of tree water and its potential sources, mainly the different soil layers, is a powerful method to infer root water uptake, while causing minimal disturbances to both the trees and the soil (Ceperley et al. 2024, Sprenger et al. 2016). During root water uptake, typically, no isotopic fractionation occurs, therefore the isotopic composition of the water that flows through the xylem vessels should reflect that of its sources (Dawson and Ehleringer 1991). Apparent isotopic fractionation resulting from methodological artifacts can bias inference on water uptake from isotopic analyses (Barbeta et al. 2022, Barbeta et al. 2019, Chen et al. 2020), but these apparent biases are consistent within species and can be corrected (Duvert et al. 2022). In temperate and beech forests, analyses of water isotopes have shown that trees would increase the proportion of water taken up from deeper soil layers as topsoil water decreases (Brinkmann et al. 2019, Geßler et al. 2022). These findings suggest that beech water uptake could exhibit certain plasticity in response to drought. However, although this plasticity could contribute to resist transient drought periods, in the mid- to long-term, reduced topsoil moisture and water uptake from this layer could compromise nutrient uptake (Querejeta, Ren and Prieto 2021). Nutrients concentrate in the upper soil (Jobbágy and Jackson 2001) as do ECM fungi (Neville et al. 2002). Thus, although reductions in topsoil water could be compensated with shifts in water uptake towards deep soil layers, reduced water uptake from the topsoil might impact negatively on nutrient uptake and on the activity and functionality of the ECM fungal community.

Here, we assess how seasonal and geographic variations in water uptake are linked to changes in the ECM fungal community on a geographic gradient along the southernmost distribution limit of *F. sylvatica*. We expected greater decoupling between water uptake and diversity and colonization by ECM fungi at sites with lower water availability, and in the driest part of the growing season. Our aims were: (*i*) to assess changes in seasonal water uptake patterns; (*ii*) to characterise the seasonal and geographical variability of the ECM fungal communities and, (*iii*) to link changes in ECM fungal colonization and community structure to water uptake patterns.

## Material and methods

### Study sites

We conducted our study in the north of Spain, where European beech reaches its southern distribution limit (except for some small isolated populations in Sicily –Italy– and central and coastal Spain, Leuschner 2020). In this area, we selected three mature monospecific beech forests along a latitudinal gradient (Table 1, Figure S1). The climate in the three selected sites varied markedly (Table 1 and Figure S1). The northernmost and rainiest site, Artikutza (Navarra, Spain, hereafter “Northern site”), was the closest to the coast and experienced a temperate oceanic climate with a mean annual precipitation of 2488.1±62.1 mm and a mean annual temperature of 15.5±1.2℃ (mean±se, 1970-2015, according to the closest meteorological station of the Regional Basque Meteorological Agency, Euskalmet). The second site, Iturrieta (Álava, Spain, hereafter “Intermediate site”), experienced a predominantly Atlantic climate with a mean annual precipitation of 1151.4±64.4 mm and mean annual temperature of 13.3±1.5℃ (2001-2015, Euskalmet). The southernmost and most continental site, Diustes (Soria, Spain, hereafter “Southern site”) experienced a Mediterranean climate with a mean annual precipitation of 779.4±20.5 mm and mean annual temperature of 14.1±1.6℃ (1967-2017, Spanish Bureau of Meteorology). To assess the impact of regional and intra-annual climatic variability on the ECM fungal community, besides climate, we prioritised management history and homogeneity in soil properties as our selection criteria, because these are crucial drivers of the ECM fungal community composition and structure (Correia et al. 2021, Rodriguez-Uña et al. 2019, Rodríguez-Uña et al. 2024). Our selected sites had been free of major impacts for ∼90 years (except for some minor and occasional wood extraction), had a non-calcareous bedrock and were located on acidic soils (pH < 5.5, Table 1). Time since the end of major activities was checked with dendrochronological analyses of wood cores extracted from all selected trees (see *Experimental design*). Wood cores were extracted in August 2020 using a Pressler increment borer (Haglöf, Långsele, Sweden). The wood cores were air-dried, glued onto wooden mounts, polished until tree rings were visible and then visually cross-dated (Stokes and Smiley 1968). Soil geology was identified on maps of the Spanish Geographic Institute and pH was measured on a set of soil samples from various depths, collected at each site.

**Table 1.**
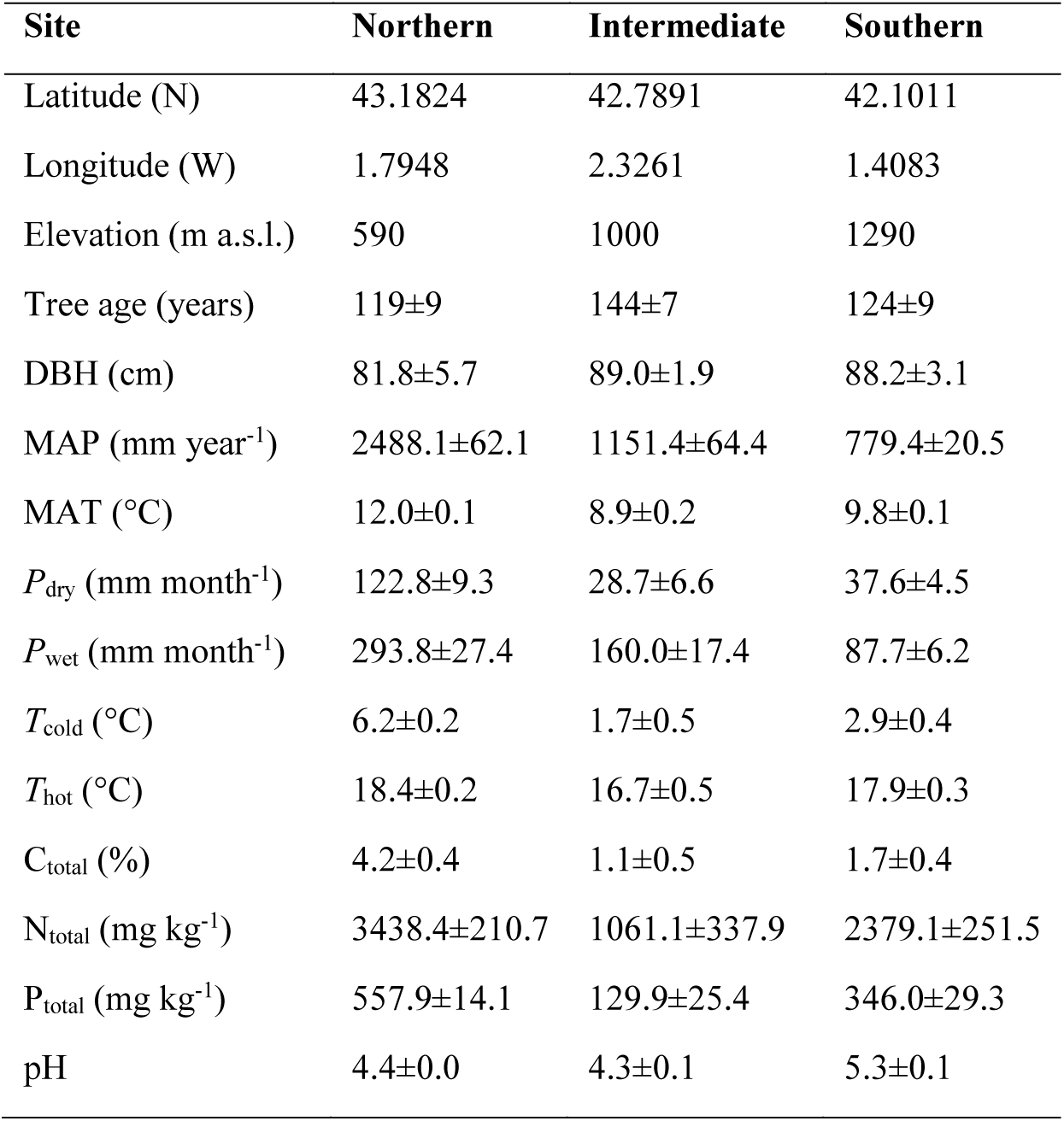
Location, mean (±se) age and size (diameter at 1.3 m height, DBH) of sampled trees, mean climatic values and mean soil properties of the three study sites in northern Spain: Northern, (Artikutza, Navarra), Intermediate (Iturrieta, Álava) and Southern (Diustes, Soria). Climatic values are: mean annual precipitation (MAP) and temperature (MAT), mean monthly precipitation of the driest (*P*_dry_) and wettest (*P*_wet_) months, mean monthly temperature of the coldest (*T*_cold_) and hottest (*T*_hot_) months. Soil properties are: total carbon (C) and nitrogen (N) contents and pH of the upper soil (0-5 cm). Climatic data provided by the Basque Regional Meteorological Agency (Euskalmet, for the Northern and Intermediate sites) and by the Spanish Meteorological Agency (AEMET, for the Southern site).

### Experimental design and sample collection

At each study site, we chose eight mature (>90 years) dominant individuals of *F. sylvatica* (3 sites × 8 trees/site = 24 trees in total). We performed two sampling campaigns, one in June 2020 (spring campaign), when we did not expect water availability to be limiting, and a second one in August 2020 (summer campaign), when water availability was expected to limit the physiological activity of both the vegetation and the associated fungal community. The same selected trees were sampled in both campaigns.

For analyses of ECM fungal diversity and to determine colonization of fine roots by ECM fungi, on each campaign and at each tree, we selected four points on the forest floor located 1-1.5 m from the base of the tree. In June, these points were located in north, south, west and east directions and in August, we selected four similar points in northwest, northeast, southeast and southwest direction. At each one of these sampling points, we dug a shallow (0-15 cm) hole (∼15 cm in diameter) and collected fine root segments. Fine root segments were transferred to Ziploc® bags with ∼200 g of accompanying soil. These bags were transported in a cool-box to the laboratory, where they were stored at 4°C until they were processed, within the following 48 h. In total, we collected 192 samples of fine roots (3 sites × 8 trees/site × 4 samples/tree × 2 campaigns = 192 samples). In addition, at one of the four locations per tree and season, we also collected an intact 5-10 cm root segment of diameter > 2 mm in each campaign, to determine fine root density per unit of soil volume (see below).

Samples for analyses of water isotopic composition and volumetric water content (VWC) were collected from five out of the eight selected trees at each study site and on each campaign. The same five trees per site were sampled in both campaigns. We collected one lignified branch segment (diameter: 1-2 cm, length: ∼10 cm) per tree and for those trees where branches were not accessible, we collected a small wood core (∼10 cm) extracted at 1.3 m height. Bark, phloem and cambium were removed before placing the branch samples or small wood cores into Exetainers® glass vials, sealed with Parafilm®. Soil samples for analyses of soil VWC and water isotopic composition were collected from a location 1.5-2 m away from the base of each tree and at least 1 m away from any of the sampling points selected for fine root collection (to avoid sampling some disturbed topsoil that might have undergone some evaporative enrichment). At each selected location, we dug a hole down to 50 cm and collected soil samples at three depth intervals: 0-5, 20-25 and 40-50 cm. All samples were collected into screw-cap glass vials and sealed with Parafilm®. In addition, at each study site and on each sampling campaign, we collected groundwater from local fountains for analyses of its isotopic composition. All samples for isotopic analyses were placed in a cool-box and once in the laboratory, they were stored at -20°C. In total, we collected 30 branches or cores (3 sites × 5 trees/site × 2 campaigns = 30 samples), 90 soil samples (3 sites × 5 trees/site × 3 depths × 2 campaigns = 90 samples) and 6 water samples (3 sites × 2 campaigns = 6 samples) for analyses of water isotopic composition.

### Characterization of soil physical and chemical properties

We estimated VWC from gravimetric water content (*W*_b_, obtained from the difference between wet and dry masses of the soil samples used for isotopic analyses, see below) using bulk density pedotransfer functions, including organic carbon and texture (Martin, Reyes and Taguas 2017). Soil chemical and physical (texture) properties were measured from additional soil samples collected in November 2020. For this latter sample collection, we randomly selected three locations at each site among the sampled trees. At each location, we dug a 50 cm hole and collected soil samples at the same depth intervals as for water isotopic analyses (0-5, 20-25 and 40-45 cm), i.e. we collected a total of 27 samples (3 sites × 3 locations/site × 3 depths/location = 27 samples). These soil samples were sieved (2 mm) and oven dried (48 h at 70°C) for texture, pH and nutrient content analyses. Texture analyses involved estimating percentages of sand, silt and clay by laser diffraction. Nutrient analyses consisted of the determination of total carbon, nitrogen and phosphorous (by UV-spectroscopy and following Kjeldhal digestion); [NH_4+_], [NO_3-_] (by UV-spectroscopy) and [K^+^] (by flame spectroscopy); pH and conductivity.

### Cryogenic water extraction and analyses of water isotopic composition

Water extraction from branches or cores and soil samples was performed by cryogenic vacuum distillation using the design and methodology proposed by (Orlowski et al. 2013), following the methodology detailed in (Jones et al. 2017). Briefly, the cryogenic extraction line consisted of six branches each with four vacuum extraction lines connected on one end to the glass vials containing the samples and on the other end to U-shaped tubes. At the onset of the extraction, samples in the glass vials were quickly frozen by immersing them into liquid N. The extraction line was then evacuated down to an atmospheric (static) pressure <1 Pa, and then the U-shape tubes were immersed in liquid nitrogen to create a cold trap. At the same time, samples were immersed in a water bath at ambient temperature, the water bath was gradually heated up to 80°C for 1 h and samples remained in the heated bath at 80°C for 2 h (i.e. total extraction time: 3 h). Pressure in the extraction line was continuously monitored with sub-atmospheric pressure sensors (APG100 Active Pirani Vacuum Gauges, Edwards, Burgess Hill, UK) to check that the lines remained leak-tight throughout the entire extraction and that the water extraction had ended. Samples were weighed before and after the extraction and before and after being oven-dried for 24 h at 105°C to assess the water extraction efficiency and to calculate sample relative water content. The extraction efficiency was always above 98%, ensuring that the isotopic composition of extracted water had not been affected by incomplete Rayleigh distillation processes (Araguasaraguas et al. 1995, Ceperley et al. 2024).

We measured the isotopic composition (δ^2^H and δ^18^O) of the extracted water samples with an off-axis integrated cavity optical spectrometer (TIWA-45EP, Los Gatos Research, USA) coupled to a liquid auto-sampler and vaporiser (LC-xt, PAL systems, Switzerland). All measured values were calibrated using two internal standards and expressed on the VSMOW-SLAP scale (Jones et al. 2017). When analysing water samples extracted from plant tissues with laser-based instruments, the presence of organic compounds (ethanol and methanol mainly) can lead to large isotopic biases (Martín-Gómez et al. 2015). Therefore, we developed a post-correction algorithm, specific for our instrument, to correct for the presence of organic compounds based on the narrowband (for methanol) and broadband (for ethanol) metrics of the absorption spectra (Leen et al. 2011).

Recent studies have shown that cryogenically extracted water from plant samples does not always reflect the isotopic composition of its sources, particularly for δ^2^H (Barbeta et al. 2020, Chen et al. 2020, de la Casa et al. 2021, Zhao et al. 2016). Barbeta et al. (2022) demonstrated that this would be caused by xylem sap water (the water that runs through the conduits connecting the roots with the leaves) being more enriched than cryogenically extracted bulk xylem water. This is because bulk xylem water consists of a mix of xylem sap water and storage water and only the former reflects source water, whereas the latter is isotopically depleted with respect to the sources (Barbeta et al. 2022). Here, we estimated the isotopic composition of xylem sap water (δ_sap_, the one that reflects the source) from bulk xylem water (δ_tree-bulk_, cryogenically extracted) by subtracting to our measurements of δ_tree-bulk_ the mean difference between δ_tree-bulk_ and δ_sap_. We calculated this mean difference from a dataset obtained from a separate set of samples of the same species (*F. sylvatica*). This set of samples consisted of 35 branch samples collected on seven dates in 2019 from five adult beech trees, similar in age and height to ours, at the Ciron Valley (France, see Barbeta et al. 2019, for the site description). Each branch sample was divided in two sub-samples, one used for extraction of δ_sap_, using the centrifugation method (Barbeta et al. 2022), and a second used for extraction of δ_tree-bulk_, using the same cryogenic distillation line described above. Mean (±sd) differences in isotopic composition between sap and bulk water (δ_tree-bulk_ - δ_sap_) were: -13.8±0.83 and -0.65±0.19‰, for δ^2^H and δ^18^O, respectively (*n* = 35), and there were no significant differences neither among trees, nor among dates (Martín-Gómez, unpublished data).

In certain soil types, estimates of plant water uptake based on bulk soil water (δ_soil-bulk_) can be biased, as they assume that plants can access tightly bound soil water (Duvert et al. 2022). This tightly bound water has been shown to be isotopically enriched, whereas unbound water available for plant uptake would be isotopically depleted with respect to bulk soil water (Chen et al. 2021). In sandy soils and/or poor in organic matter, the proportion of tightly bound organic matter is negligible, but this is not the case for very clayey soils and/or rich in organic matter. At our forest sites, the soils were relatively poor in organic matter (Table S1), but the soils at two of the sites had very high clay content (>20%). Hence, for these two sites (Northern and Southern), we estimated the isotopic composition of unbound soil water (δ_soil-u_) as follows:

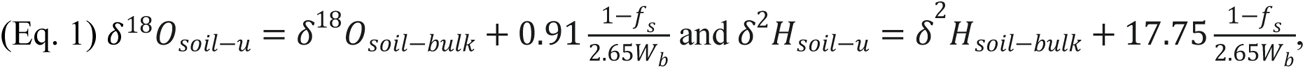

where *f*_s_ is the sand fraction (mean for each soil layer calculated from soil profiles dug at three locations per site, see previous section) and *W*_b_ is the gravimetric water content of a given soil sample, calculated for the same samples from the difference in weight before cryogenic water extraction (wet mass, *M*_w_) and after being oven dried (48 h at 105°C, dry mass, *M*_D_), according to: *W*_b_ = (*M*_w_ *M*_D_)/*M*_D_. These formulations (Eq. 1) are based on empirical measurements and consider surface fractionation effects that occur within the soil matrix (Chen, Auerswald and Schnyder 2016). Henceforth, isotopic composition of soil water refers to corrected unbound soil water for the clayey sites (Northern and Southern) and to bulk soil water for the sandier site (Intermediate).

We used measurements of water isotopic composition to determine the relative contribution to corrected tree xylem sap water (δ_tree-sap_) of each one of the potential water sources using Bayesian mixing models (Parnell et al. 2010). These models assume that all considered water sources can be accessed simultaneously by the plants. To produce plausible simulations of the most likely relative contribution of each source the models use Markov Chain Monte Carlo (MCMC) methods (Stock, Jackson, Ward, Parnell, Phillips and Semmens 2018). Our model inputs were mean and standard deviation of water isotopic compositions (δ^2^H and δ^18^O) of soil water. We used estimated unbound soil water (δ_soil-u_) at the Northern and Southern sites, with very clayey soils, and bulk soil water (δ_soil-bulk_) at the Intermediate site. Mean and standard deviation of isotopic compositions were calculated for each source per study site and sampling campaign. Soil water from intermediate (20-25 cm) and deep (40-45 cm) soil layers were pooled together because their isotopic compositions were not statistically different (Figure S2). The target values were individual isotopic compositions of each tree and sampling campaign. Trophic enrichment factor was set to 0, as we assumed that no fractionation occurred during root water uptake. We also assumed no concentration dependency. We ran 400,000 iterations and discarded the first 200,000. Convergence was checked according to the Gelman and Geweke diagnostics. All models were run in R version 4.1.3 (Team 2021) using the *MixSIAR* package (Stock and Semmens 2016).

### Fine root density and measurements of colonization by ECM fungi

Root samples were carefully washed in the laboratory and for each sample (bag with fine roots and accompanying soil collected at a unique location), we randomly selected three fine root (<2 mm) segments of 5 cm each. Root colonization by ECM fungi was assessed by counting the number of branched ECM root tips in each segment under the stereomicroscope, always by the same two observers. We then estimated ECM fungal colonization per cm of fine root for each location as the average of the three segments.

The additional larger root segment (diameter >2 mm, one segment collected per tree and sampling campaign) was rinsed and scanned with WinRHIZO® (Regent Instruments Inc., Québec, Canada). From this analysis, we used the total length of fine roots per unit of soil volume (in cm m^-3^) to calculate a standardized rate of ECM fungal colonization as the number of live ECM tips per cubic meter of soil.

### Molecular identification of ECM fungi

All live ECM tips counted from the three randomly selected segments from each sample were collected into Eppendorf tubes, i.e. we generated one sample (one Eppendorf tube) per sampled location (that is four samples per tree and campaign). DNA was extracted from these mycorrhizal root tips using the DNeasy PowerSoil Pro Kit (Qiagen, CA, USA) following manufacturer’s instructions. Extraction negative controls were also created to ensure lack of contamination during DNA extraction. The following steps of molecular analysis and bioinformatics were carried out at a commercial sequencing facility (www.allgenetics.eu). First, a fragment of the internal transcribed spacer (ITS) genomic region of the fungal rRNA (ca 400 bp) was amplified using the primers ITS1-F (5’ CTT GGT CAT TTA GAG GAA GTA A 3’) (Gardes and Bruns 1993) and ITS2 (5’ GCT GCG TTC TTC ATC GAT GC 3’) (White et al. 1990). PCRs were carried out in a final volume of 12.5 μL, containing 1.25 μL of template DNA, 0.5 μM of the primers, 3.25 μL of Supreme NZYTaq 2x Green Master Mix (NZYTech), and ultrapure water up to 12.5 μL. The reaction mixture was incubated as follows: an initial denaturation step at 95°C for 5 min, followed by 30 cycles of 95°C for 30 s, 47.7°C for 45 s, 72°C for 45 s, and a final extension step at 72°C for 7 min. The libraries were run on 2% agarose gels stained with GreenSafe (NZYTech) and imaged under UV light to verify the library size. A second PCR round (with identical conditions but only 5 cycles and 60°C as the annealing temperature) was carried out to attach the oligonucleotide indices required for multiplexing different libraries in the same sequencing pool. Subsequent library purification was performed using the Mag-Bind RXNPure Plus magnetic beads (Omega Biotek). Then, libraries were pooled in equimolar amounts according to the quantification data provided by the Qubit dsDNA HSAssay (Thermo Fisher Scientific). The pool was sequenced on an Illumina MiSeq platform (Illumina, San Diego, CA, USA) using PE300 v3 kits.

The obtained ITS1 amplicon reads were processed using DADA2 (Callahan, McMurdie, Rosen, Han, Johnson and Holmes 2016), implemented in QIIME 2 (release 2021.2) (Bolyen, Rideout, Dillon, Bokulich, Abnet, Al-Ghalith, Alexander, Alm, Arumugam and Asnicar 2019). PCR primers were removed, reads were denoised, merged and filtered according to their quality, removing the chimaeric reads and clustering the resulting sequences into Amplicon Sequence Variants (ASVs).

Taxonomy was assigned to the obtained 2273 ASVs with a pre-trained classifier of the UNITE reference database (Abarenkov et al. 2020, updated in February 2020), using the feature-classifier classify-sklearn approach implemented in QIIME2 (Bokulich et al. 2018). We discarded 534 ASVs (23%) that corresponded to (i) singletons (i.e., ASVs containing only one member sequence in the whole data set), (ii) ASVs occurring at a frequency below 0.01% in each sample, or (iii) unidentified sequences and sequences assigned only at the kingdom level (‘Fungi’). Based on the alpha rarefaction curves created with the number of ASVs obtained for each sample, six samples were discarded because they did not reach the plateau in the number of ASVs observed. ECM fungal communities were characterized with the 675 ASVs (40%, from a total of 1688 ASVs) identified as ECM fungi at the genus or species level, also including 78 ASVs corresponding to families only composed by ECM fungi (i.e. Boletaceae, Clavulinaceae, Elaphomycetaceae, Russulaceae and Thelephoraceae). We manually matched the 183 ASVs that reached the genus level with the UNITE reference database to improve its taxonomy assignment, reaching a total of 209 and 466 ASVs identified at the genus and species level, respectively (based on a threshold similarity >97%). For simplicity, hereafter we refer to ASVs as “species”, although we acknowledge that ASVs do not necessarily correspond to actual species.

### Diversity and richness indexes, community composition and functional diversity for ECM fungi

Calculations of ECM fungal community structure, diversity and richness were based on data of presence-absence of species in each sample. We calculated our indexes for individual trees from the four samples collected per tree and campaign. Species abundances were calculated for each tree and campaign based on the presence of a given species measured in each one of the four samples per tree and campaign. We calculated the Shannon-Wiener index (*H’*) using the *diversity* function from R package *vegan* (Oksanen et al. 2019) and according to:

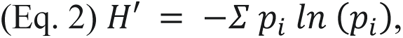

where *p*_*i*_ is the relative abundance of species *i* per tree calculated as the number of records found for species *i* (ranging from 0 to 4 for each species and tree) divided by the total number of records found for each tree. Species richness was estimated as the total number of species present in each sample (*S*), using the *specnumber* function from R package *vegan* and according to:

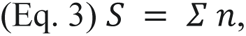

Where *n* is the number of species in each of the four samples of each tree and campaign.

Community composition for each tree and campaign was characterized with the identity and relative abundance of each species (*p*_*i*_). As a proxy for functional diversity of the ECM fungal community, we classified species according to their capacity to explore the surrounding soil based on their hyphal development, the so-called “exploration type”, with two categories: long-exploration (including long- and medium-exploration types) vs. short-exploration (including short-exploration and contact mycelium, Agerer 2001). Each species was assigned to one of these two categories following (Agerer 2006, Defrenne et al. 2019, Ostonen et al. 2017). We found 9 species out of a total of 151, for which the exploration type was unknown and therefore these were excluded for further calculations. We calculated the proportion (*χ*) of species of each functional type (long, *χ*_*l*_, vs. short, *χ*_*s*_) for each sample, according to:

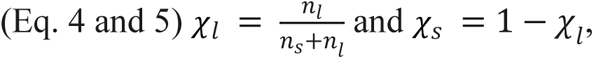

where *n_l_* and *n_s_* indicate the number of species of each type. We calculated a value of *χ*_*l*_ and *χ*_*s*_ for each sample, hence our database contained four values per tree and sampling campaign.

### Statistical analyses

All calculations and analyses were run in R version 4.1.3 (R Team 2021). Differences in soil physical and chemical properties among study sites (Northern, Intermediate and Southern) and soil depths (0-5, 25-30 and 45-50 cm) were assessed by means of analysis of variance (ANOVA). For the rest of our variables, we assessed differences among study sites, between sampling campaigns (June, spring, and August, summer), among sampling depths (for soil VWC only) and their interactions, using linear mixed models (LMMs, for soil VWC, ECM fungal colonization, total fine root length per unit of soil volume, number of ECM live tips per m^3^ of soil and Shannon-Wiener diversity index) or generalized linear mixed models (GLMMs, with a Poisson error distribution for species richness, and with a binomial error distribution for proportion of ECM functional group, long- vs. short exploration type), including tree as a random factor. Differences in tree and soil water isotopic composition between sampling campaigns and among depths (for the soil) were also assessed with LMMs, including tree as random factor, but not study site as a fixed factor. Instead, we ran separate LMMs for each study site because we were not interested in differences across study sites due to geographic variability in isotopic composition of precipitation water (Bowen, Wassenaar and Hobson 2005). In addition, we tested for the potential effects of shallow (0-5 cm) soil VWC (measured at individual trees on each study site and on each sampling campaign) on ECM fungal species richness and diversity using LMMs. We fitted all LMMs and GLMMs using packages *lme4* (Bates, Maechler, Bolker and Walker 2015) and *lmerTest* (Kuznetsova, Brockhoff and Christensen 2017) to estimate *P*-values. For all models with significant results for the effect of study site, post-hoc tests were run using *emmeans* (Lenth et al. 2020). Graphs to visualize our results were generated with *ggplot2* (Wickham, Chang and Wickham 2016).

We assessed differences among study sites and sampling campaigns on ECM fungal community composition with a permutational multivariate analysis of variance (PERMANOVA), using the *adonis* function with 9,999 permutations. The analysis was based on Bray-Curtis because it enables the estimation of community composition dissimilarities considering relative abundances of species. To visualize differences in ECM fungal community composition among study sites and sampling campaigns, we used an ordination analysis of the communities with a non-metric multidimensional scaling (NMDS), based on the Bray-Curtis similarity index, using function *metaMDS* (all functions from R package *vegan*, Oksanen et al. 2019).

## Results

### Soil volumetric water content, chemical and physical properties

Soil physical and chemical properties were homogenous within sites, although there were some differences for certain properties (Table S1). Most notably, we found that the soils were sandier at the Intermediate site (Iturrieta). In addition, the soil was richer (in terms of total C, N and P content) at the Northern site (Artikutza) and less acidic at the Southern site (Diustes, Table S1). Soil volumetric water content (VWC) was higher in spring (June) than in summer (August) and the upper soil (0-5 cm) was always wetter than the other soil layers (20-25 and 40-45 cm), at all sites (Figure 1). There were also some significant differences in VWC among study sites: the Intermediate site (Iturrieta) had lower mean VWC than the Northern (Artikutza) and Southern (Diustes) sites (Figure 1, Table 2). The site with the highest mean annual precipitation (Northern, Artikutza) did not have the highest VWC, instead the site with the highest VWC was the Southern one (Diustes).

**Figure 1.**
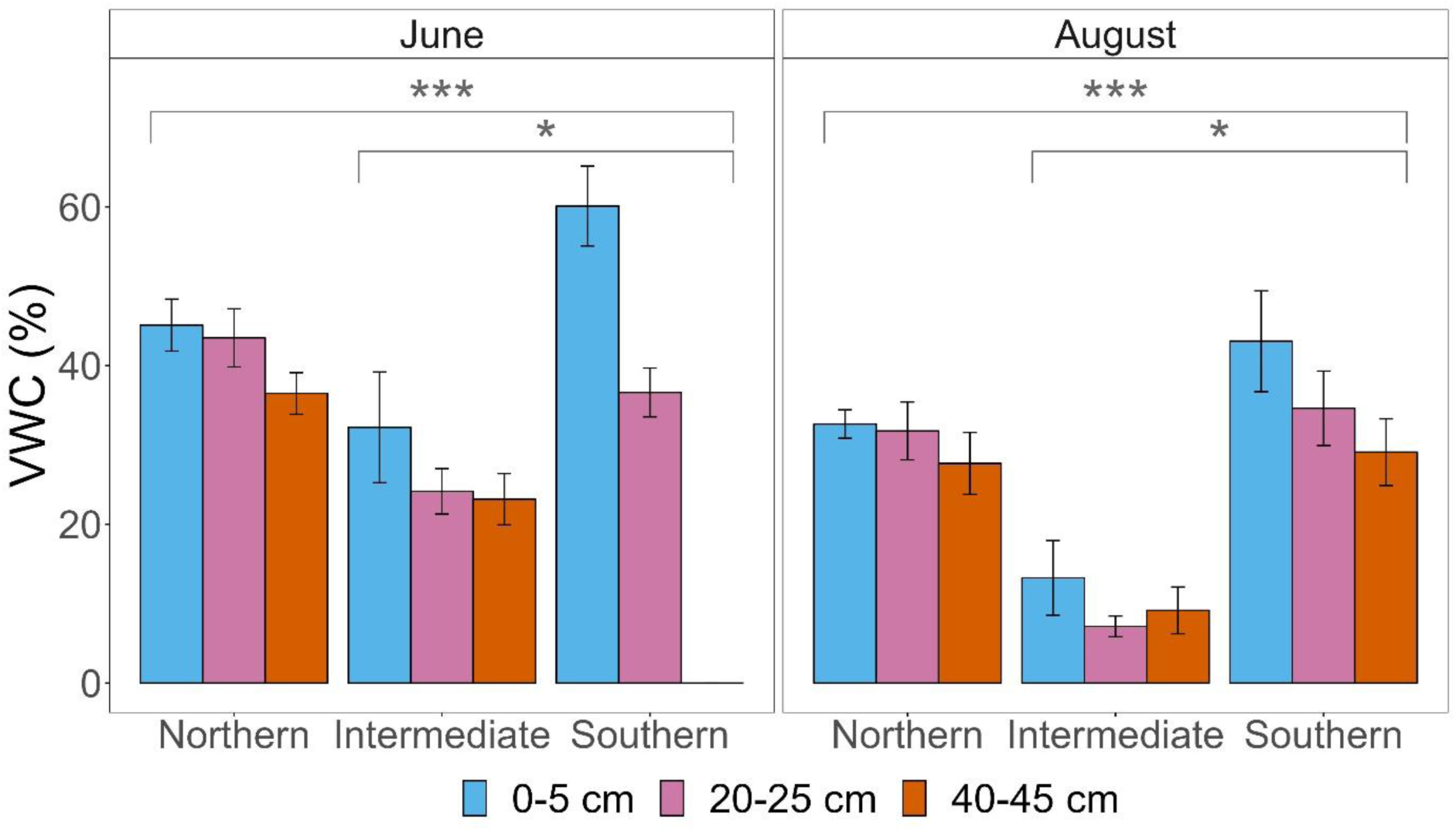
Mean (±se, *n* = 5) soil volumetric water content (VWC) for different soil depths at the three study sites (Northern, Intermediate and Southern) and during the two sampling campaigns (spring, June, and summer, August; no soil samples were collected at 40-45 cm depth at the Southern site in June). Asterisks indicate significant differences between sites.

**Table 2.** Results of the linear or generalised mixed models for differences among sites (Northern, Intermediate and Southern), campaigns (spring, and summer), depths (0-5 cm and 20-50 cm) and their interactions in: VWC; root density; ECM fungal colonization, richness, diversity and fraction of short exploration type. Marginal (*R^2^_m_*) and conditional (*R^2^_c_*) coefficients indicate variance explained by fixed (*R^2^_m_*) or fixed and random (*R^2^_c_*) factors.

| **Soil Volumetric Water Content (VWC)** |  |  |  |  |  |
| --- | --- | --- | --- | --- | --- |
| **Source** | **Df** | **F** | **P** | **$R^2_m$** | **$R^2_c$** |
| Site | 2 | 27.22 | <0.001 | 0.66 | 0.69 |
| Campaign | 1 | 34.4 | <0.001 |  |  |
| Soil depth | 2 | 10.61 | <0.001 |  |  |
| Site × Campaign | 2 | 1.36 | 0.265 |  |  |
| Site × Depth | 4 | 2.23 | 0.077 |  |  |
| Campaign × Depth | 2 | 1.22 | 0.304 |  |  |
| **Fine root density (log-transformed)** |  |  |  |  |  |
| **Source** | **Df** | **F** | **P** | **$R^2_m$** | **$R^2_c$** |
| Site | 2 | 2.58 | 0.099 | 0.17 | 0.24 |
| Campaign | 1 | 3.83 | 0.064 |  |  |
| Site × Campaign | 2 | 0.49 | 0.617 |  |  |
| **Colonization (log-transformed)** |  |  |  |  |  |
| **Source** | **Df** | **F** | **P** | **$R^2_m$** | **$R^2_c$** |
| Site | 2 | 9.04 | 0.001 | 0.26 | 0.27 |
| Campaign | 1 | 44.92 | <0.001 |  |  |
| Site × Campaign | 2 | 0.40 | 0.674 |  |  |
| **Density of live tips (log-transformed)** |  |  |  |  |  |
| **Source** | **Df** | **F** | **P** | **$R^2_m$** | **$R^2_c$** |
| Site | 2 | 7.57 | <0.001 | 0.49 | 0.58 |
| Campaign | 1 | 32.54 | <0.001 |  |  |
| Site × Campaign | 2 | 0.12 | 0.124 |  |  |
| **Species richness (Poisson distribution)** |  |  |  |  |  |
| **Source** | **Df** | **$\chi^2$** | **P** | **$R^2_m$** | **$R^2_c$** |
| Site | 2 | 4.05 | 0.137 | 0.06 | 0.25 |
| Campaign | 1 | 0.10 | 0.749 |  |  |
| Site × Campaign | 2 | 4.00 | 0.136 |  |  |
| **Species diversity (Shannon Index)** |  |  |  |  |  |
| **Source** | **Df** | **F** | **P** | **$R^2_m$** | **$R^2_c$** |
| Site | 2 | 3.07 | 0.067 | 0.13 | 0.13 |
| Campaign | 1 | 0.26 | 0.613 |  |  |
| Site × Campaign | 2 | 0.29 | 0.752 |  |  |
| **Short exploration type (Binomial distribution)** |  |  |  |  |  |
| **Source** | **Df** | **$\chi^2$** | **P** | **$R^2_m$** | **$R^2_c$** |
| Site | 2 | 9.57 | 0.008 | 0.08 | 0.08 |
| Campaign | 1 | 0.92 | 0.337 |  |  |
| Site × Campaign | 2 | 0.48 | 0.787 |  |  |

### Water isotopic composition of plant water and its sources

We found that there were some isotopic offsets between plant water and its potential sources (Figure S2). When we applied our corrections and estimated xylem sap water (δ_sap_) and unbound soil water (δ_soil-u_, for the clayey sites, Northern and Southern) these offsets were minimised. After these corrections, the estimated values of δ_sap_ overlapped with the isotopic composition of the potential sources in the dual isotopic space (Figure S2). At the sites with high clay content, this overlap became more evident when we incorporated the correction for estimated unbound soil water (δ_soil-u_, Figure S2).

The isotopic compositions of soil water from 40-50 and 20-25 cm depths were not statistically different, neither for δ^18^O nor for δ^2^H, regardless of the study site or sampling campaign (Figure S3). Thus, values from 40-45 and 20-25 cm were pooled together for subsequent analyses. The δ^18^O of the topsoil (0-5 cm) was more enriched than that of the deeper soil layers (20-25 and 40-45 cm, Table 3), suggesting isotopic enrichment of the topsoil. Also, the interaction term (campaign × depth) was significant at the Northern and Intermediate sites (Table 3). This implies that the isotopic differentiation along the soil profile was stronger in summer, when evaporative enrichment of the topsoil would have been stronger, than in spring. In contrast, δ^2^H of the upper soil water was not significantly enriched with respect to that of deeper soil layers, likely because δ^2^H is less affected by evaporative enrichment than δ^18^O due to the lower atomic mass of H. The δ^18^O and δ^2^H of the soil varied between campaigns at all sites (Table 2, Figure 2). Meanwhile, δ^18^O and δ^2^H of estimated tree sap water did not differ between campaigns, in any of the sites (Table 3; Figure 2).

**Figure 2.**
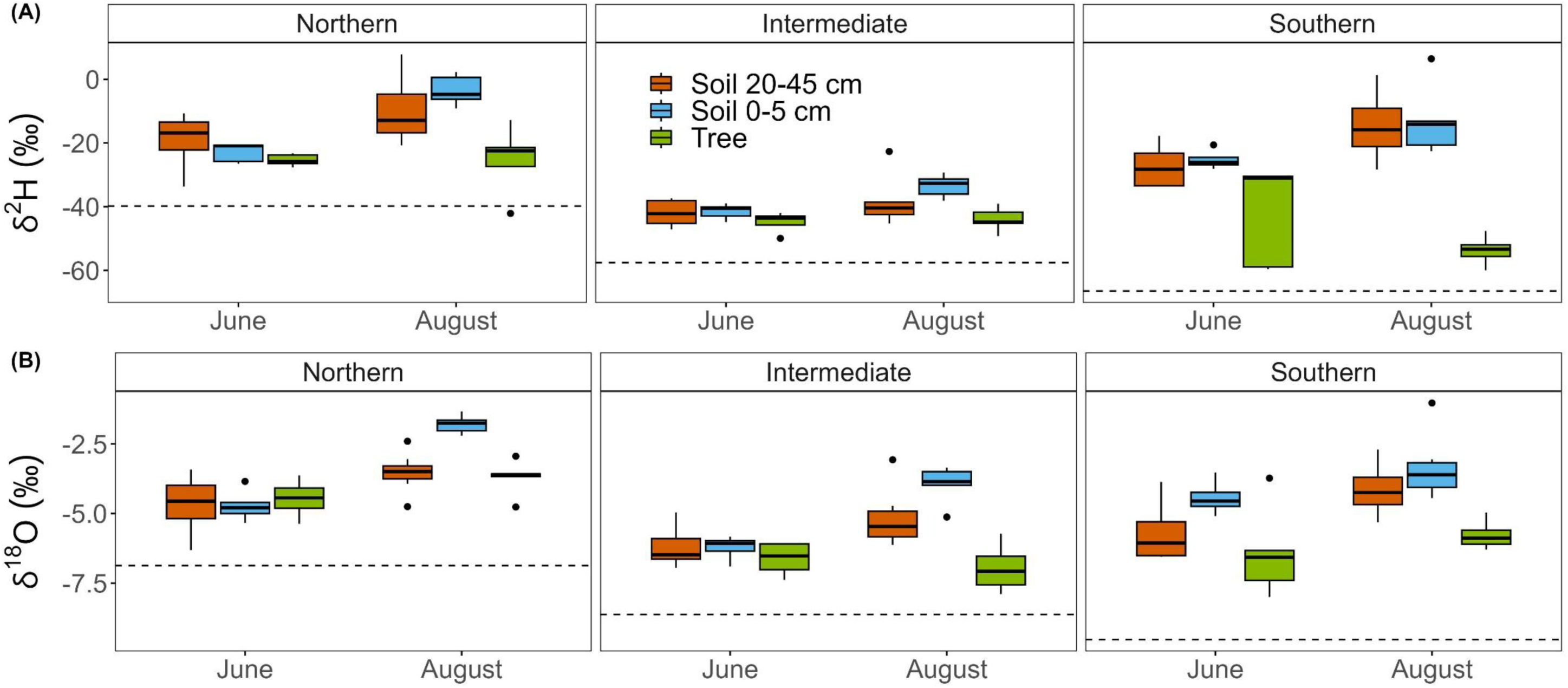
Boxplots of the hydrogen (δ^2^H, A) and oxygen (δ^18^O, B) stable isotope composition for estimated tree sap water (green boxes) and of estimated available soil water for 0-5 cm (blue boxes) and 20-45 cm (orange boxes), for the three sites (Northern, Intermediate and Southern) and the two campaigns (spring, June, and summer, August). Estimated δ^2^H and δ^18^O of soil water are that of unbound soil water for the clayey sites (Northern and Southern) and that of bulk soil water for the sandy site (Intermediate). Dashed lines depict δ^2^H and δ^18^O of groundwater at each site.

**Table 3.** Results of the linear mixed models to test for differences in soil and tree water stable isotope composition (δ^2^H and δ^18^O) between sampling campaigns (spring, June, and summer, August), sampling depths (for soil water only) and their interactions, at each one of the sites (Northern, Intermediate and Southern). Tree water δ^2^H and δ^18^O are those of estimated sap water and soil water δ^2^H and δ^18^O are those of estimated available soil water: unbound soil water for the clayey sites (Northern and Southern) and bulk soil water for the sandier site (Intermediate). Marginal (*R^2^_m_*) and conditional (*R^2^_c_*) coefficients indicate the variance explained by fixed only and by fixed and random factors, respectively. Df: degrees of freedom. Statistically significant *P*-values are shown in bold.

| Variable | Source | Northern |  |  |  |  | Intermediate |  |  |  |  | Southern |  |  |  |  |
| --- | --- | --- | --- | --- | --- | --- | --- | --- | --- | --- | --- | --- | --- | --- | --- | --- |
| | | Df | $F$ | $P$ | $R^2_m$ | $R^2_c$ | Df | $F$ | $P$ | $R^2_m$ | $R^2_c$ | Df | $F$ | $P$ | $R^2_m$ | $R^2_c$ |
| Soil $\delta^2\text{H}$ | Campaign | 1 | 34.74 | **<0.001** | 0.44 | 0.68 | 1 | 8.72 | **0.008** | 0.29 | 0.48 | 1 | 20.71 | **<0.001** | 0.32 | 0.62 |
|  | Soil depth | 1 | 0.16 | 0.707 |  |  | 1 | 3.04 | 0.097 |  |  | 1 | 0.4 | 0.534 |  |  |
| | Campaign × Depth | 1 | 4.92 | **0.037** | | | 1 | 2.57 | 0.125 | | | 1 | 0.02 | 0.894 | | |
| Soil $\delta^{18}\text{O}$ | Campaign | 1 | 84.74 | **<0.001** | 0.69 | 0.83 | 1 | 37.71 | **<0.001** | 0.56 | 0.71 | 1 | 17.96 | **<0.001** | 0.42 | 0.63 |
|  | Soil depth | 1 | 17.63 | **<0.001** |  |  | 1 | 5.19 | **0.035** |  |  | 1 | 10.44 | **0.005** |  |  |
| | Campaign × Depth | 1 | 18.66 | **<0.001** | | | 1 | 6.46 | **0.02** | | | 1 | 0.27 | 0.609 | | |
| Tree $\delta^2\text{H}$ | Campaign | 1 | <0.01 | 0.975 | <0.01 | <0.01 | 1 | 0.15 | 0.722 | 0.02 | 0.03 | 1 | 3.17 | 0.149 | 0.24 | 0.26 |
| Tree $\delta^{18}\text{O}$ | Campaign | 1 | 3.22 | 0.147 | 0.26 | 0.26 | 1 | 0.54 | 0.504 | 0.06 | 0.06 | 1 | 0.8 | 0.422 | 0.07 | 0.07 |

The results of the MixSIAR model predicted that there was a seasonal shift in the relative contributions of topsoil and deeper soil water pools at the Northern and Intermediate site, but not at the Southern site (Figure 3). At the Southern site, the estimated relative contribution of the topsoil (0-5 cm) was low (Figure 3), despite high water availability in this soil layer (Figure 1). At the Southern site, water uptake relied ∼90% on deep soil, both in spring (89±9%, estimated ± sd) and summer (92±13%, Figure 3). In contrast, at the Northern site, estimated percentages of water uptake varied markedly between spring and summer (66±19 and 34±19% in spring and 14±11 and 86±11% in summer, for topsoil and deep soil, respectively, Figure 3). At the Intermediate site, estimated water uptake in spring was evenly distributed between topsoil and deep soil (45±23 and 55±23%, respectively), whereas in the summer the estimated contribution of the topsoil decreased to 19±11% (Figure 3).

**Figure 3.**
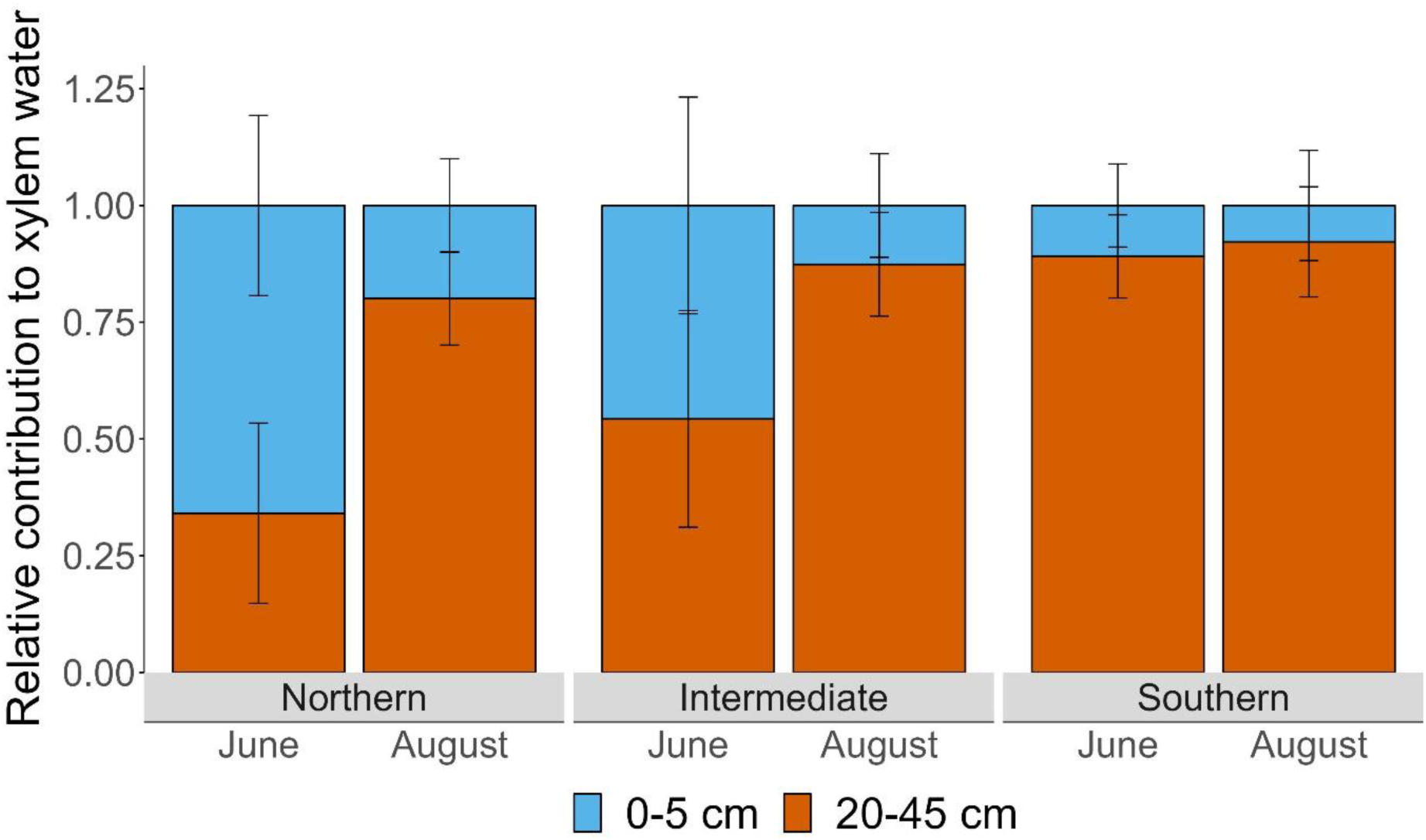
Relative contribution to tree xylem sap water of topsoil (0-5 cm) and deep (20-25 and 40-15 cm) soil water, for the three study sites (Northern, Intermediate and Southern) and the two sampling campaigns (spring, June, and summer, August). Bars are the mean (±sd) estimated relative contribution according to Bayesian mixing models.

### Fine root density and ECM fungal colonization

Mean (± se) fine root density (total length of fine roots per unit of soil volume) was marginally larger (Table 2) in spring (June, 561±73 cm m^-3^) than in summer (August, 399±39 cm m^-3^). Among sites, fine root density was marginally larger at the Southern site (593±96 cm m^-3^) than at the Northern site (382±58 cm m^-3^), whereas density of fine roots at the Intermediate site (464±56 cm m^-3^) was not different from either site. Colonization of fine roots by ECM fungi (number of live tips per cm of fine root) was higher in spring (June) than in summer (August), at all sites (Figure 4a, Table 2). In addition, we found more colonization of fine roots by ECM fungi at the Southern site than at the Northern and Intermediate sites (Figure 4a, Table 2). When we analysed the estimated number of live ECM tips per unit of soil volume (a variable combining number of live tips per unit of fine root length and density of fine roots), we found that there were marked differences between seasons and among sites (Table 2): density of live ECM tips per unit of soil volume dropped significantly from spring (1519±230 tips m^-3^) to summer (541±71 tips m^-3^) and was higher at the Southern site (1568±323 tips m^-3^) than at the Northern (634±147 tips m^-3^) and Intermediate sites (887±157 tips m^-3^).

**Figure 4.**
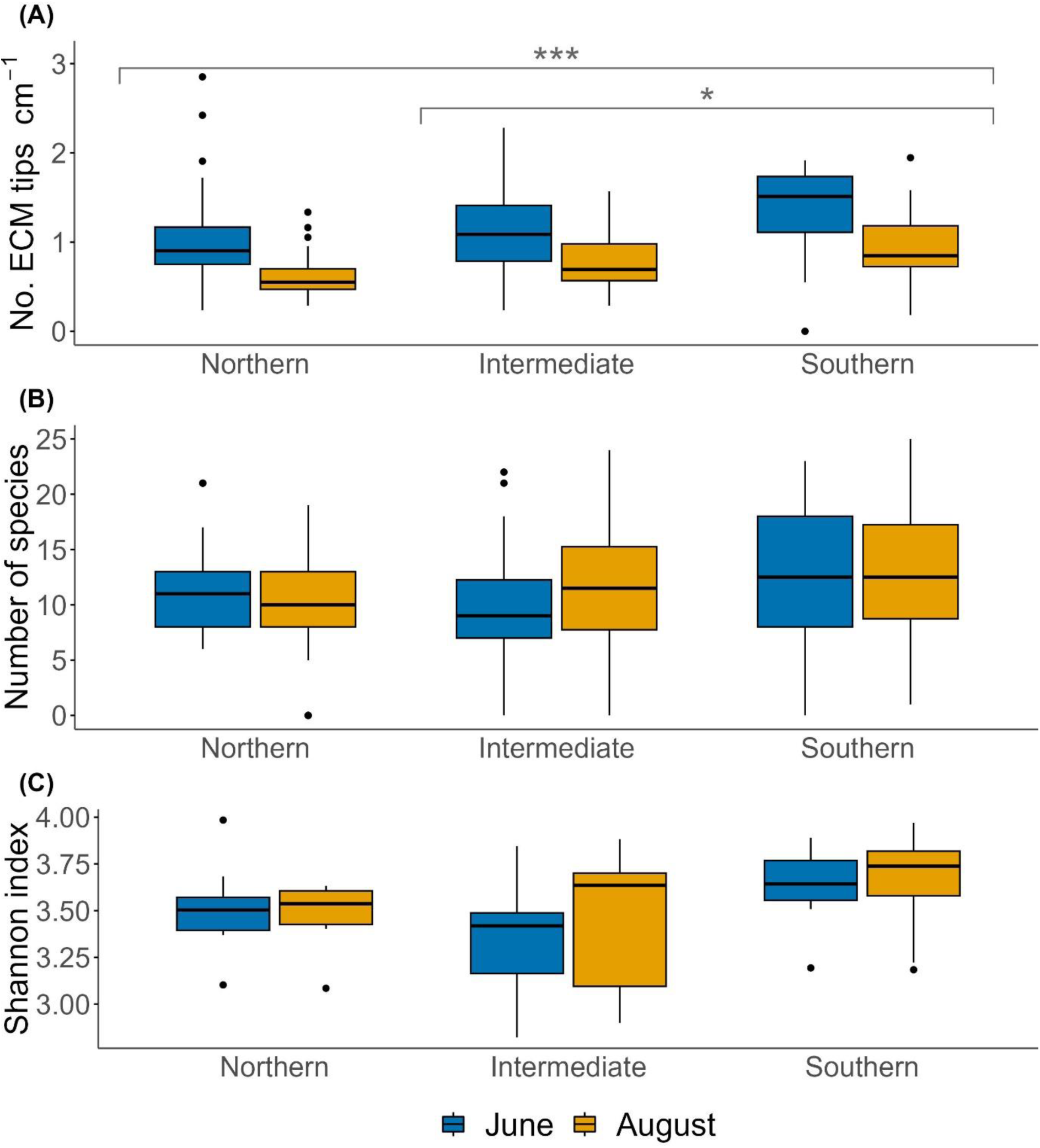
Boxplots of the ectomycorrhizal (ECM) fungal (A) colonization rate, (B) species richness and (C) species diversity for the three study sites (Northern, Intermediate and Southern) and the two sampling campaigns (spring, June, and summer, August). Asterisks indicate significant differences (\**P* < 0.05, \*\*\**P* < 0.001) between study sites, according to post-hoc pairwise comparisons. Colonization rates were higher (*P* < 0.001) in June than in August in the three sites.

### ECM fungal communities

We identified 675 ECM fungal species, of which 578 (85.6%) belonged to the phylum Basidiomycota, 93 (13.8%) to Ascomycota and 4 (0.6%) to Mucoromycota. Among these, we found 38 genera, with *Tomentella*, *Russula* and *Lactarius* being the most abundant. The number of unique species detected was 51, distributed evenly across sites (Table S2). The mean (±se) species richness per tree were 11.16 ± 0.69, 10.13 ± 0.86 and 12.47 ± 1.17 in spring and 10.19 ± 0.83, 11.47 ± 0.98 and 12.56 ± 1.07 in summer at the Northern, Intermediate and Southern sites, respectively. The species diversity per tree was 3.51 ± 0.09, 3.33 ± 0.12 and 3.63 ± 0.08 in spring and 3.48 ± 0.06, 3.46 ± 0.14 and 3.66 ± 0.11 in summer at the Northern, Intermediate and Southern sites, respectively. Our GLMM revealed no differences in ECM fungal species richness among study sites or between sampling campaigns (Figure 4b; Table 2). We found marginally significant differences among study sites on ECM fungal species diversity, with higher species diversity at the Southern site than at the Intermediate site (Figure 4c; Table 2). In addition, we found that there was greater variability for both species richness and diversity index at the Southern site (Figure 4b-c). The PERMANOVA model showed strong evidence of an effect of study site on ECM fungal community composition (Pseudo-*F* = 4.88, *P* < 0.001), but not of sampling campaign (Pseudo-*F* = 0.69, *P* = 0.97; Table S3). NMDS visualization was consistent with the effect of the study site (Figure 5).

**Figure 5.**
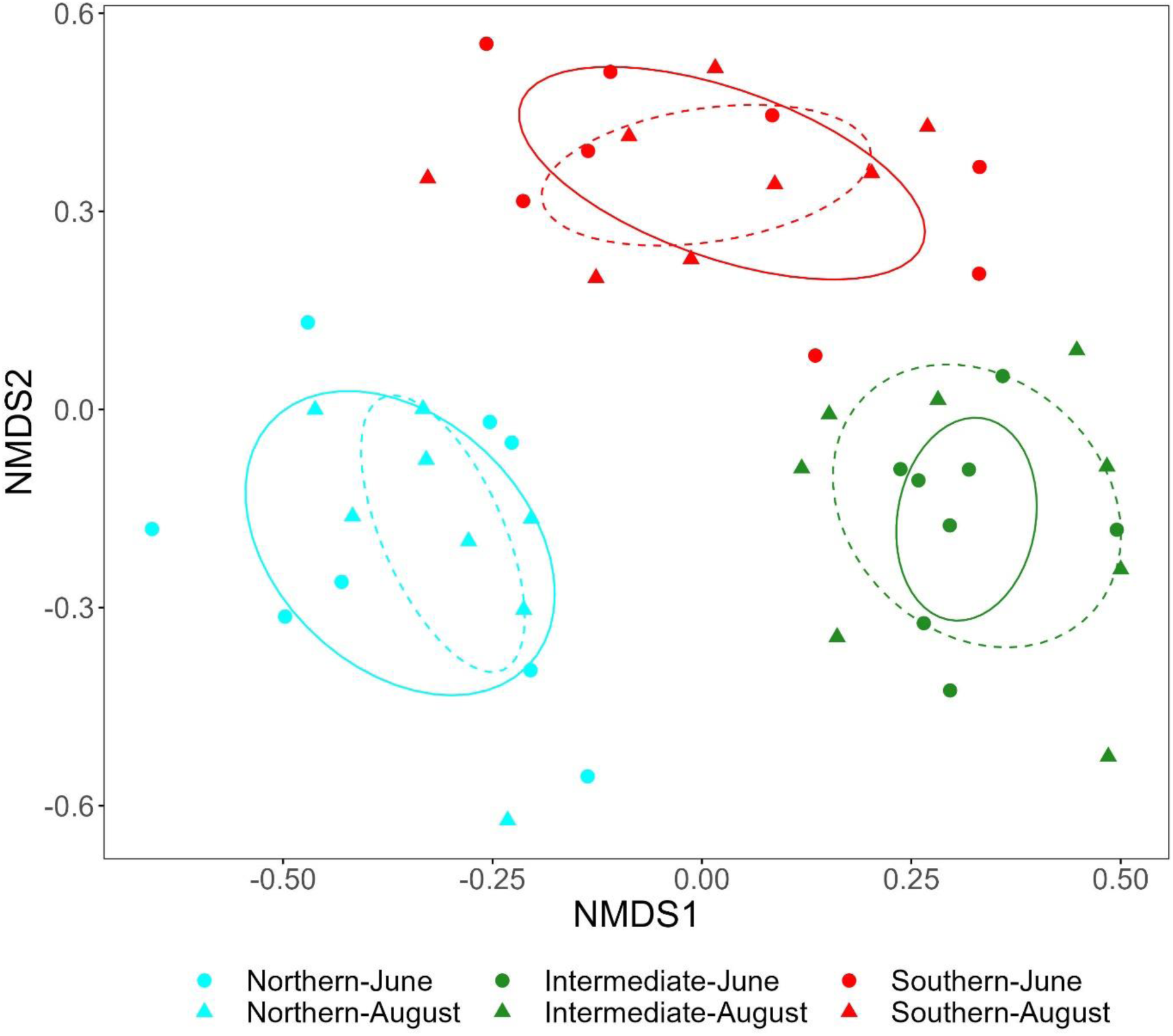
Non-metric multidimensional scaling (NMDS) plot of the ectomycorrhizal fungal communities for the three study sites (Northern, Intermediate and Southern) and the two sampling campaigns. Points and triangles on the ordination space represent June (spring) and August (summer) sampled communities, respectively, based on Bray-Curtis dissimilarity indices. Solid and dotted lines show the standard deviation ellipses of June and August sampled communities, respectively.

There were differences in the relative proportion of short- and long-exploration types of ECM fungi among sites, but not between seasons (Table 1): the Intermediate site presented the highest proportion of species with short-exploration types (and the subsequent lowest proportion of long-exploration types), being significantly different from that at the Southern site (Figure 6; Table 2). Finally, we found that VWC had a marginally significant and positive effect on species diversity, irrespective of the season, but not on species richness (Table S4).

**Figure 6.**
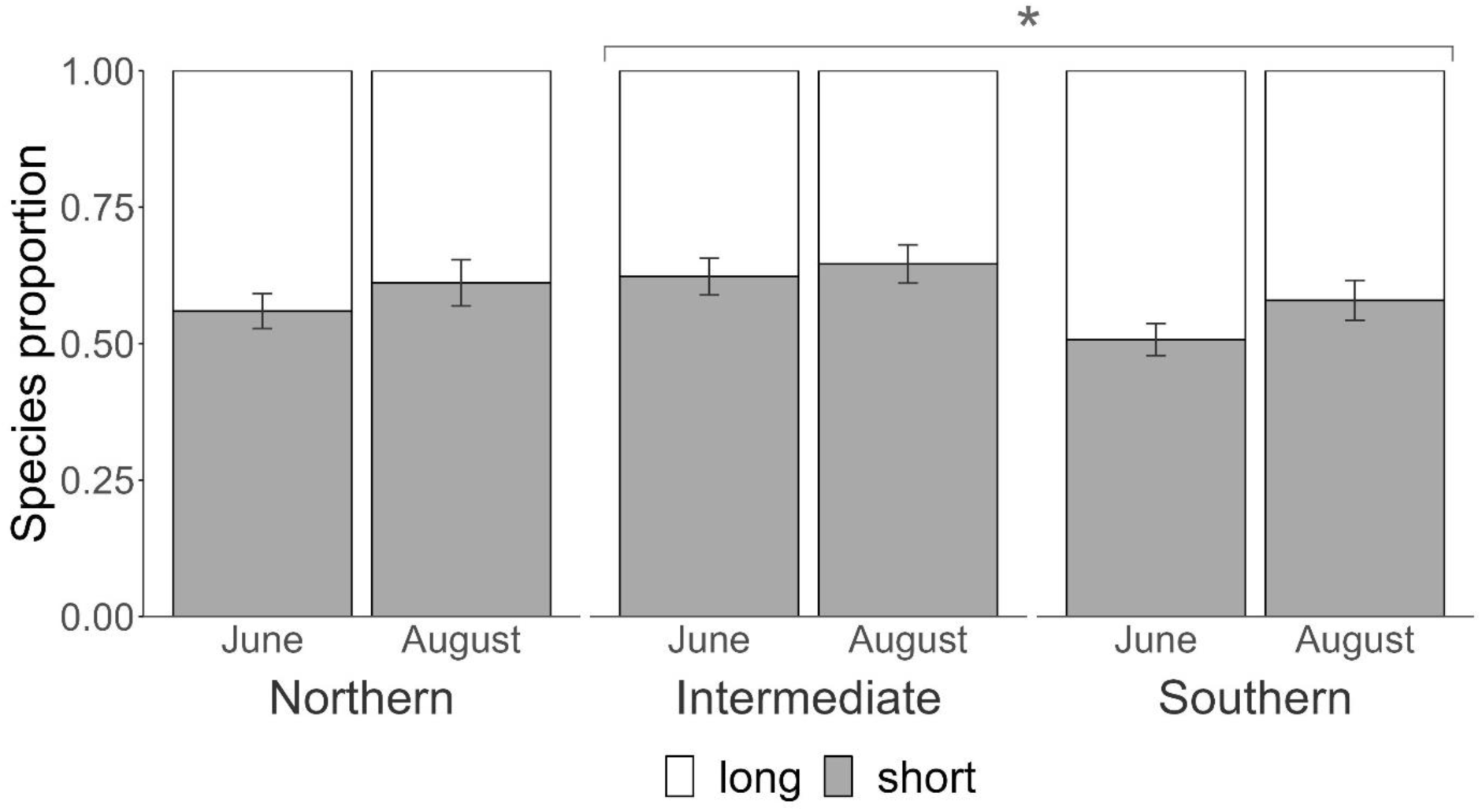
Proportion of ectomycorrhizal fungal species classified according to the exploration type of their hyphae: short-exploration vs. long-exploration. Bars depict mean (+se, *n* = 8 trees) proportions for the three study sites (Northern, Intermediate and Southern) and the two sampling campaigns (spring, June, and summer, August). Asterisk indicates significant difference (*P* < 0.05) between sites according to post-hoc pairwise comparisons.

## Discussion

We combined molecular analyses of ECM fungal community composition with patterns of root water uptake, from geographically and climatically contrasting populations of European beech. We had anticipated that seasonal and geographic differences in water availability would underlie changes in the diversity and colonization by ECM fungi as well as in the relative water uptake patterns. Our results showed greater diversity and colonization by ECM fungi when the topsoil was more humid, but the contribution to water uptake of the topsoil did not always change in response to water availability in this soil layer.

The natural distribution of European beech (*F. sylvatica*) is mainly limited by water availability (Geßler et al. 2007) and there is ample evidence showing that drought impacts negatively on the growth and health of these forests (e.g. Neycken et al. 2024, Zimmermann et al. 2015). Yet, *F. sylvatica* can depict some physiological and morphological adaptations that allow this tree to cope with moderate drought. These include, for example, a tight stomatal control in response to both atmospheric (i.e. vapour pressure deficit) and soil drought (Jonard et al. 2011, Leuschner, Schipka and Backes 2022), or reduction of the leaf to sapwood area (Arend et al. 2022). Plasticity in water uptake is another mechanism that would allow beech trees to overcome such drought periods by shifting water uptake from shallower to deeper soil layers (Kinzinger et al. 2023). Here, we found an increase in relative water uptake from deeper soil layers at the Northern and Intermediate sites in summer, similar to previous observations in other beech forests (Brinkmann et al. 2021). However, shifts in relative water uptake towards deeper soil layers may not be sufficient to compensate for reduced topsoil water availability, since greater relative water uptake does not necessary imply an increase in the absolute amount of water taken up (Geßler et al. 2022). Here, we did not quantify transpiration (the most direct proxy of tree water uptake), thus we cannot conclude whether the observed shifts in relative water uptake patterns were compensatory. Interestingly, at our Southernmost and most marginal study site, we did not find evidence of any shifts in relative water uptake between seasons. At this site, topsoil moisture was consistently high, maybe due to an evener distribution of precipitation events during the summer in comparison to, for example, our intermediate site (Figure S1); yet the estimated contribution of the topsoil to water uptake at this site was always the lowest.

We expected patterns of water uptake and of ECM fungal diversity and colonization to be coupled, to a certain extent. Our results suggest that topsoil water availability would drive both water uptake patterns and ECM fungal diversity and colonization, but with different implications. In spring, the topsoil contributed significantly to water uptake at the Intermediate and Northern sites, consistent with the higher density of fine roots found in this soil layer in this season. ECM fungal colonization was also greater in spring at all sites. Given that nutrient uptake depends on soil water availability (Schlesinger et al. 2016) and that ECM hyphae are sensitive to desiccation (Querejeta, Egerton-Warburton and Allen 2009), our results suggest that during the early part of the growing season, nutrient and water uptake would be coupled, at least in these sites. In contrast, in summer at all sites and at the Southern site also in spring, we found that topsoil contribution to water uptake was small, despite the slightly higher VWC of this layer. In summer, water uptake relied on deeper soil layers, while nutrients and ECM (although less abundant than in spring) were still concentrated in the upper soil. This could imply that under moderate drought, water uptake from deep soil layers could still sustain the transpiration demand, but nutrient uptake could be compromised (Querejeta, Ren and Prieto 2021). It could be argued that hydraulic lift could partially alleviate this potential limitation, but intraspecific hydraulic lift in beech is limited (Hafner, Hesse and Grams 2021, Zapater et al. 2011). Thus, under increasing drought, although trees could compensate for reduced topsoil water availability by shifting water uptake towards deeper soil layers, the associated ECM fungal community dwelling in the upper soil would still suffer from desiccation.

Beyond seasonal differences, across our geographic gradient, we found that our study sites not only differed in ECM fungal colonization, but also in community structure. Analyses of simple metrics (species richness and diversity indexes) did not shed light on how ECM fungal communities varied across the sampled geographic gradient, nor between seasons. We only found that the ECM fungal community was marginally more diverse at the Southernmost site. In contrast, analyses of community composition based on species relative abundances showed that ECM fungal communities clearly differed in their structure across sites but not between seasons within sites. Our forests did not differ in tree species composition (monospecific stands) nor in land-use history (mature forests). Thus, our results indicate that the prevailing climate, more than the seasonal variability, together with other topographic and abiotic conditions (soil properties, slope, etc.), determined how ECM fungal communities structured (e.g. Miyamoto et al. 2015).

We found a greater proportion of long-exploration ECM types at the Southernmost and most marginal site. At the Intermediate site, which had a higher sand fraction and the lowest soil VWC, the proportion of short-exploration types was more abundant. Experiments with saplings and under controlled conditions have shown that under drought, the proportion of contact-type ECM fungi decreases (Castaño et al. 2023, Köhler et al. 2018). In the field, Nickel et al. (2018) also found a rapid decrease of contact-, short- and medium-exploration types in response to drought, and in the longer term (3 years of drought exposure) also a relative increase in long-exploration types. In contrast, Castaño et al. (2018) found that short-exploration types were more abundant under drier conditions while long-exploration types were more abundant under wetter and cooler conditions, in a Mediterranean forest. Similarly, Fernandez et al. (2023) found an increase in contact- and short-exploration types under reduced rainfall in boreal trees. Contact- and short-exploration types are less C-demanding; therefore, these exploration types would be favoured when photosynthesis becomes limiting (Fernandez et al. 2020), for example in sites experiencing periods with low soil water availability (Castaño et al. 2018). This could be the case of our Intermediate site, where the combination of sparse precipitation events during the summer months (Figure S1), together with the lower water holding capacity of the soil, would result in periods of low water availability that could have favoured the selection of ECM fungal partners that would be less costly to replace. In contrast, at the Southern site, we found a larger proportion of long-exploration types. Long-exploration ECM fungi usually deploy long hydrophobic hyphae that should allow for a more efficient water transport and for the exploration of larger soil volumes (Lehto and Zwiazek 2011, Prieto et al. 2016). The deployment of such structures would be advantageous in environments where water and host photosynthesis would not be limiting, as they are more costly to maintain (Fernandez et al. 2017). This could be the case at our Southern site, where we found the highest water availability, both in spring and summer, and a larger proportion of long-exploration types. At this site, the deployment of an ECM network with longer emanating exploration tissues could have also been one of the factors that might have contributed to the persistence of this marginal beech population outside its core distribution range. In the long-term and under a climate change scenario, ECM fungal communities with a higher proportion of long-exploration ECM fungi might be able to maintain their functionality to a greater extent than those dominated by contact- or short-type ECM fungi, provided drought did not impair host photosynthesis. Alternatively, under a climate change scenario with increasing frequency and severity of extreme droughts, hosting ECM networks with greater functional diversity (i.e. a mix of long- and short-exploration types) could be a more advantageous strategy conferring greater resistance and recovery capacity.

## Conclusions

We found that fine root density and colonization by ECM fungi were highest at our Southernmost and most isolated beech forest, where we also found a higher proportion of long-exploration ECM fungal types. Our results agree with previous studies on European beech suggesting that marginal populations or those at the rear-edge of their climatic distribution limit might not necessarily be the most vulnerable to the impacts of climate change (Bert et al. 2022, Cavin and Jump 2017, Vilà-Cabrera and Jump 2019). Instead, our results suggest that beyond their marginality and mean climatic values (e.g. MAT or MAP), maintaining high soil water availability throughout the growth season is more relevant for buffering the impacts of increasing drought. In contrast to previous studies, our approach is not based on the assessment of a single forest function (e.g. growth). Instead, we combined methodologies from plant ecophysiology and molecular biology to assess relative water uptake and the biodiversity of soil dwelling microorganisms, forest properties that depend on tree roots and their associations with symbiotic fungi. Our results suggest that under a climate change scenario, European beech forests could face transient reductions in topsoil water by shifting water uptake towards deeper soil layers, but these reductions would still be detrimental for the survival of their ECM fungal partners. In a scenario with increased frequency, duration and intensity of drought episodes, the ECM fungal community might progressively contract and become less diverse, compromising future productivity and nutrient cycling at the ecosystem level, even at forest sites traditionally not considered vulnerable to drought stress.

## Data availability

Illumina next-generation DNA sequences will be published in GenBank®. The rest of the data associated with this publication will be made publicly available on figshare, upon acceptance of this manuscript for publication.

## Acknowledgments

Funding was provided by the regional Government of the Basque Country (projects for basic and applied research: PIBA-2019-105) awarded to TEG and DMM and by the Spanish Ministry of Science through Grant PHLISCO (PID2019-107817RB-I00) and a Ramón y Cajal Fellowship (RYC2021-031759-I) awarded to TEG. DMM and ARU were partially supported by the Spanish State Research Agency through María de Maeztu Excellence Unit accreditation 2018-2022 (Ref. MDM-2017-0714). LW has received funding from the European Research Council (ERC) under the European Union’s Horizon 2020 research and innovation programme (Grant agreement No. 101003125) and the French Government in the framework of the IdEX Bordeaux University “Investments for the Future” program / GPR Bordeaux Plant Sciences. Special thanks to Nicolas Devert and Sabrina Dubois for the sample processing and analyses of water isotopic composition. Many thanks also to Mario Blanco-Sánchez and Marina Ramos-Muños for their assistance during DNA extraction, and to Jesús L. Angulo for his advice with statistical analyses. Thanks to Paula Martín-Gómez for providing the data to correct our tree water isotopic composition.

