## Supplementary Material for "Changes in the functional diversity and abundance of ectomycorrhizal fungi are decoupled from water uptake patterns in European beech forests"

**Supporting Information**

**Contents**

Tables S1-S4

Figures S1-S3

References

**Table S1.** Mean (±se, $n$ = 3) characteristics of the three soil layers analysed (0-5, 25-30 and 45-50 cm) at each one of the study sites. Different letters indicate significant differences among sites (capital letters A-C), among soil depths (capital letters X-Z) or among study sites and soil depths (small case letters).

| Variable | Northern (Artikutza) | | | Intermediate (Iturrieta) | | | Southern (Diustes) | | |
|---|---|---|---|---|---|---|---|---|---|
| | **0-5 cm** | **20-25 cm** | **40-45 cm** | **0-5 cm** | **20-25 cm** | **40-45 cm** | **0-5 cm** | **20-25 cm** | **40-45 cm** |
| Clay (%) | 23.1±1.9$^{A}$ | 23.7±8.3$^{A}$ | 20.8±7.1$^{A}$ | 2.7±0.5$^{B}$ | 3.7±0.1$^{B}$ | 5.0±1.2$^{B}$ | 20.4±4.2$^{A}$ | 28.5±12.1$^{A}$ | 23.5±2.5$^{A}$ |
| Loam (%) | 51.0±3.0$^{A}$ | 49.6±3.0$^{A}$ | 51.2±0.4$^{A}$ | 24.4±1.9$^{B}$ | 22.7±2.3$^{B}$ | 27.0±2.9$^{B}$ | 56.5±3.0$^{A}$ | 52.8±7.7$^{A}$ | 57.6±3.6$^{A}$ |
| Sand (%) | 26.0±2.1$^{A}$ | 26.7±5.9$^{A}$ | 28.0±7.0$^{A}$ | 72.9±2.4$^{B}$ | 73.6±3.3$^{B}$ | 68.0±4.1$^{B}$ | 23.0±2.2$^{A}$ | 18.7±4.43$^{A}$ | 18.8±2.7$^{A}$ |
| $C_{total}$ (%) | 5.4±0.2$^{AX}$ | 3.5±0.2$^{AY}$ | 3.3±0.2$^{AY}$ | 2.9±1.2$^{BX}$ | 0.3±0.1$^{BY}$ | 0.3±0.1$^{BY}$ | 2.9±0.3$^{BX}$ | 1.6±0.5$^{BY}$ | 0.6±0.3$^{BY}$ |
| $N_{total}$ (g $kg^{-1}$) | 3.9±0.2$^{AX}$ | 3.3±0.3$^{AY}$ | 3.0±0.4$^{AY}$ | 2.2±0.5$^{BX}$ | 0.490±0.1$^{BY}$ | 0.4±0.12$^{BY}$ | 3.3±0.1$^{CX}$ | 2.2±0.1$^{CY}$ | 1.6±0.1$^{CY}$ |
| $P_{total}$ (g $kg^{-1}$) | 0.6±0.0$^{AX}$ | 0.5±0.0$^{AX}$ | 0.5±0.0$^{AX}$ | 0.2±0.0$^{BY}$ | 0.1±0.0$^{BY}$ | 0.1±0.0$^{BY}$ | 0.4±0.0$^{CY}$ | 0.3±0.0$^{CY}$ | 0.3±0.0$^{CY}$ |
| K (mg $kg^{-1}$) | 20.7±3.6$^{abcde}$ | 9.8±2$^{de}$ | 7±1.7$^{e}$ | 25.1±0.9$^{ab}$ | 34.1±2.3$^{cde}$ | 57.1±7.5$^{bcde}$ | 43.7±11.7$^{abcd}$ | 11.5±3.1$^{abc}$ | 16.5±8$^{a}$ |
| $NO_3^-$ (mg $kg^{-1}$) | 1.2±0.3 | 0.6±0.3 | 0.7±0.2 | 3.2±1.2 | 1.1±0.4 | 1.1±0.3 | 0.7±0.3 | 0.9±0.3 | 0.8±0.3 |
| $NH_4^+$ (mg $kg^{-1}$) | 4.8±1.8$^{cdefgh}$ | 4.3±0.6$^{bcdefgh}$ | 3.4±0.4$^{efghi}$ | 13.3±3.4$^{a}$ | 2.2±0.3$^{fghi}$ | 2±0.3$^{hi}$ | 3.6±0.5$^{defgh}$ | 2.1±0.1$^{ghi}$ | 1.2±0.1$^{i}$ |
| Conductivity (µS $cm^{-1}$) | 106.1±13.4$^{X}$ | 79.6±8.4$^{Y}$ | 68±2.9$^{Y}$ | 158.4±40.2$^{X}$ | 56.5±8.7$^{Y}$ | 55.5±10.3$^{Y}$ | 132.9±8.3$^{X}$ | 75.6±8.5$^{Y}$ | 50.6±5.2$^{Y}$ |
| pH | 4.3±0$^{B}$ | 4.5±0$^{B}$ | 4.5±0$^{B}$ | 4.2±0.1$^{B}$ | 4.2±0.1$^{B}$ | 4.4±0.1$^{B}$ | 5.4±0.2$^{A}$ | 5.3±0.2$^{A}$ | 5.3±0.1$^{A}$ |

**Table S2.** Taxonomical classification of the ectomycorrhizal fungal species identified among sites and sampling campaigns (J = June, spring; A = August, summer), and their corresponding mycelium exploration type.

| **Phylum** | **Class** | **Order** | **Family** | **Genus** | **Species** | **Exploration** | **Northern** | **Intermediate** | **Southern** |
|---|---|---|---|---|---|---|---|---|---|
| Basidiomycota | Agaricomycetes | Agaricales | Agaricaceae | Lepiota | *Lepiota elseae* | unknown | | A | |
| Basidiomycota | Agaricomycetes | Agaricales | Amanitaceae | Amanita | *Amanita fulva* | medium-long | | J, A | |
| Basidiomycota | Agaricomycetes | Agaricales | Amanitaceae | Amanita | *Amanita olivaceogrisea* | medium-long | | | J |
| Basidiomycota | Agaricomycetes | Agaricales | Amanitaceae | Amanita | *Amanita pantherina* | medium-long | A | | |
| Basidiomycota | Agaricomycetes | Agaricales | Amanitaceae | Amanita | *Amanita subparvipantherina* | medium-long | J, A | | |
| Basidiomycota | Agaricomycetes | Agaricales | Cortinariaceae | Cortinarius | *Cortinarius alnetorum* | medium-long | J, A | A | |
| Basidiomycota | Agaricomycetes | Agaricales | Cortinariaceae | Cortinarius | *Cortinarius decipiens* | medium-long | A | J, A | |
| Basidiomycota | Agaricomycetes | Agaricales | Cortinariaceae | Cortinarius | *Cortinarius paranomalus* | medium-long | A | | |
| Basidiomycota | Agaricomycetes | Agaricales | Cortinariaceae | Cortinarius | *Cortinarius quarciticus* | medium-long | | A | |
| Basidiomycota | Agaricomycetes | Agaricales | Cortinariaceae | Cortinarius | *Cortinarius subtilis* | medium-long | | J, A | |
| Basidiomycota | Agaricomycetes | Agaricales | Cortinariaceae | Cortinarius | *Cortinarius tacitus* | medium-long | | J | |
| Basidiomycota | Agaricomycetes | Agaricales | Cortinariaceae | Cortinarius | *Cortinarius torvus* | medium-long | | A | |
| Basidiomycota | Agaricomycetes | Agaricales | Cortinariaceae | Cortinarius | | medium-long | | J, A | |
| Basidiomycota | Agaricomycetes | Agaricales | Cortinariaceae | Cortinarius | | medium-long | | A | |
| Basidiomycota | Agaricomycetes | Agaricales | Cortinariaceae | Cortinarius | | medium-long | | J, A | |
| Basidiomycota | Agaricomycetes | Agaricales | Cortinariaceae | Cortinarius | | medium-long | | A | |
| Basidiomycota | Agaricomycetes | Agaricales | Cortinariaceae | Cortinarius | | medium-long | A | J, A | J, A |
| Basidiomycota | Agaricomycetes | Agaricales | Cortinariaceae | Cortinarius | | medium-long | J, A | J, A | J |
| Basidiomycota | Agaricomycetes | Agaricales | Cortinariaceae | Cortinarius | | medium-long | A | | J |
| Basidiomycota | Agaricomycetes | Agaricales | Cortinariaceae | Cortinarius | | medium-long | | J | |
| Basidiomycota | Agaricomycetes | Agaricales | Cortinariaceae | Cortinarius | | medium-long | J, A | J | |
| Basidiomycota | Agaricomycetes | Agaricales | Cortinariaceae | Cortinarius | | medium-long | A | | |
| Basidiomycota | Agaricomycetes | Agaricales | Cortinariaceae | Cortinarius | | medium-long | J | | |
| Basidiomycota | Agaricomycetes | Agaricales | Cortinariaceae | Cortinarius | | medium-long | J | | |
| Basidiomycota | Agaricomycetes | Agaricales | Hydnangiaceae | Laccaria | *Laccaria angustilamella* | medium-long | | | J |
| Basidiomycota | Agaricomycetes | Agaricales | Hydnangiaceae | Laccaria | | medium-long | | | J |
| Basidiomycota | Agaricomycetes | Agaricales | Hygrophoraceae | Hygrophorus | *Hygrophorus eburneus* | contact-short | A | A | J, A |
| Basidiomycota | Agaricomycetes | Agaricales | Inocybaceae | Inocybe | *Inocybe assimilata* | contact-short | J, A | J, A | J, A |
| Basidiomycota | Agaricomycetes | Agaricales | Inocybaceae | Inocybe | *Inocybe hirtella* | contact-short | | J, A | |
| Basidiomycota | Agaricomycetes | Agaricales | Inocybaceae | Inocybe | *Inocybe krieglsteineri* | contact-short | | A | |
| Basidiomycota | Agaricomycetes | Agaricales | Inocybaceae | Inocybe | *Inocybe napipes* | contact-short | J | J | A |
| Basidiomycota | Agaricomycetes | Agaricales | Inocybaceae | Inocybe | *Inocybe petiginosa* | contact-short | J, A | A | J, A |
| Basidiomycota | Agaricomycetes | Agaricales | Inocybaceae | Inocybe | *Inocybe xanthomelas* | contact-short | J | | |

| | | | | | | | | | |
|---|---|---|---|---|---|---|---|---|---|
| Basidiomycota | Agaricomycetes | Agaricales | Inocybaceae | Inocybe | unidentified | contact-short | J | J | J |
| Basidiomycota | Agaricomycetes | Agaricales | Inocybaceae | Inocybe | unidentified | contact-short | J | J | |
| Basidiomycota | Agaricomycetes | Agaricales | Tricholomataceae | Mycenella | *Mycenella bryophila* | unknown | | A | |
| Basidiomycota | Agaricomycetes | Agaricales | Tricholomataceae | Tricholoma | *Tricholoma hemisulphureum* | medium-long | | A | |
| Basidiomycota | Agaricomycetes | Atheliales | Atheliaceae | Byssocorticium | *Byssocorticium atrovirens* | contact-short | | J, A | J, A |
| Basidiomycota | Agaricomycetes | Atheliales | Atheliaceae | Byssocorticium | unidentified | contact-short | J, A | | |
| Basidiomycota | Agaricomycetes | Atheliales | Atheliaceae | Piloderma | *Piloderma lanatum* | medium-long | | A | |
| Basidiomycota | Agaricomycetes | Atheliales | Atheliaceae | Piloderma | unidentified | medium-long | A | J, A | J, A |
| Basidiomycota | Agaricomycetes | Boletales | Boletaceae | Boletus | *Boletus reticulatus* | medium-long | A | | |
| Basidiomycota | Agaricomycetes | Boletales | Boletaceae | Imleria | *Imleria badia* | unknown | J, A | A | A |
| Basidiomycota | Agaricomycetes | Boletales | Boletaceae | Leccinum | *Leccinum pseudoscabrum* | medium-long | J, A | J, A | J, A |
| Basidiomycota | Agaricomycetes | Boletales | Boletaceae | Octaviania | *Octaviania asterosperma* | medium-long | J | | |
| Basidiomycota | Agaricomycetes | Boletales | Boletaceae | Strobilomyces | *Strobilomyces strobilaceus* | unknown | | J | |
| Basidiomycota | Agaricomycetes | Boletales | Boletaceae | Tylopilus | *Tylopilus felleus* | medium-long | | | A |
| Basidiomycota | Agaricomycetes | Boletales | Boletaceae | Xerocomellus | *Xerocomellus poederi* | medium-long | J | | |
| Basidiomycota | Agaricomycetes | Boletales | Boletaceae | Xerocomellus | *Xerocomellus porosporus* | medium-long | | J, A | |
| Basidiomycota | Agaricomycetes | Boletales | Boletaceae | Xerocomellus | *Xerocomellus pruinatus* | medium-long | J | J, A | J |
| Basidiomycota | Agaricomycetes | Boletales | Boletaceae | Xerocomellus | | medium-long | | | A |
| Basidiomycota | Agaricomycetes | Boletales | Boletaceae | | | medium-long | J | J | J |
| Basidiomycota | Agaricomycetes | Boletales | Sclerodermataceae | Scleroderma | *Scleroderma citrinum* | medium-long | J, A | J, A | |
| Basidiomycota | Agaricomycetes | Cantharellales | Cantharellaceae | Craterellus | *Craterellus tubaeformis* | contact-short | J, A | J, A | J, A |
| Basidiomycota | Agaricomycetes | Cantharellales | Cantharellales_fam_ Incertae_sedis | Sistotrema | *Sistotrema oblongisporum* | medium-long | J | | |
| Basidiomycota | Agaricomycetes | Cantharellales | Cantharellales_fam_ Incertae_sedis | Sistotrema | unidentified | medium-long | J, A | J, A | J, A |
| Basidiomycota | Agaricomycetes | Cantharellales | Clavulinaceae | Clavulina | *Clavulina rugosa* | medium-long | | J, A | |
| Basidiomycota | Agaricomycetes | Cantharellales | Clavulinaceae | Clavulina | | medium-long | J, A | J, A | J, A |
| Basidiomycota | Agaricomycetes | Cantharellales | Clavulinaceae | Membranomyces | unidentified | unknown | A | J, A | J, A |
| Basidiomycota | Agaricomycetes | Cantharellales | Clavulinaceae | | | medium-long | J, A | J, A | A |
| Basidiomycota | Agaricomycetes | Cantharellales | Hydnaceae | Hydnum | *Hydnum ellipsosporum* | medium-long | J | A | A |
| Basidiomycota | Agaricomycetes | Cantharellales | Hydnaceae | Hydnum | *Hydnum subovoideisporum* | medium-long | | | J |
| Basidiomycota | Agaricomycetes | Cantharellales | Hydnaceae | Hydnum | unidentified | medium-long | J, A | J, A | J, A |
| Basidiomycota | Agaricomycetes | Cantharellales | Hydnaceae | Hydnum | | medium-long | J, A | J, A | J, A |
| Basidiomycota | Agaricomycetes | Cantharellales | Hydnaceae | Hydnum | | medium-long | J | | |
| Basidiomycota | Agaricomycetes | Gomphales | Gomphaceae | Ramaria | unidentified | medium-long | | J | |
| Basidiomycota | Agaricomycetes | Hysterangiales | Hysterangiaceae | Hysterangium | *Hysterangium nephriticum* | medium-long | | J | |
| Basidiomycota | Agaricomycetes | Hysterangiales | Hysterangiaceae | Hysterangium | *Hysterangium thwaitesii* | medium-long | | A | |
| Basidiomycota | Agaricomycetes | Russulales | Russulaceae | Lactarius | *Lactarius blennius* | contact-short | | J, A | J, A |

| | | | | | | | | | |
|---|---|---|---|---|---|---|---|---|---|
| Basidiomycota | Agaricomycetes | Russulales | Russulaceae | Lactarius | *Lactarius camphoratus* | contact-short | A | J, A | A |
| Basidiomycota | Agaricomycetes | Russulales | Russulaceae | Lactarius | *Lactarius illyricus* | contact-short | A | | |
| Basidiomycota | Agaricomycetes | Russulales | Russulaceae | Lactarius | *Lactarius pallidus* | contact-short | | J | |
| Basidiomycota | Agaricomycetes | Russulales | Russulaceae | Lactarius | *Lactarius pterosporus* | contact-short | J, A | J, A | |
| Basidiomycota | Agaricomycetes | Russulales | Russulaceae | Lactarius | *Lactarius rostratus* | contact-short | J, A | J | J, A |
| Basidiomycota | Agaricomycetes | Russulales | Russulaceae | Lactarius | *Lactarius rubrocinctus* | contact-short | A | J, A | J, A |
| Basidiomycota | Agaricomycetes | Russulales | Russulaceae | Lactarius | *Lactarius tabidus* | contact-short | J, A | | |
| Basidiomycota | Agaricomycetes | Russulales | Russulaceae | Lactarius | | contact-short | J, A | J, A | J, A |
| Basidiomycota | Agaricomycetes | Russulales | Russulaceae | Lactarius | | contact-short | | J, A | A |
| Basidiomycota | Agaricomycetes | Russulales | Russulaceae | Russula | *Russula acrifolia* | contact-short | | A | A |
| Basidiomycota | Agaricomycetes | Russulales | Russulaceae | Russula | *Russula anthracina* | contact-short | A | | |
| Basidiomycota | Agaricomycetes | Russulales | Russulaceae | Russula | *Russula brevipes* | contact-short | | J | J |
| Basidiomycota | Agaricomycetes | Russulales | Russulaceae | Russula | *Russula brunneoviolacea* | contact-short | J | A | A |
| Basidiomycota | Agaricomycetes | Russulales | Russulaceae | Russula | *Russula chloroides* | contact-short | J, A | | |
| Basidiomycota | Agaricomycetes | Russulales | Russulaceae | Russula | *Russula curtipes* | contact-short | J, A | | |
| Basidiomycota | Agaricomycetes | Russulales | Russulaceae | Russula | *Russula cyanoxantha* | contact-short | J, A | | |
| Basidiomycota | Agaricomycetes | Russulales | Russulaceae | Russula | *Russula densifolia* | contact-short | J, A | J, A | J, A |
| Basidiomycota | Agaricomycetes | Russulales | Russulaceae | Russula | *Russula fellea* | contact-short | J, A | J, A | J, A |
| Basidiomycota | Agaricomycetes | Russulales | Russulaceae | Russula | *Russula foetens* | contact-short | | J | |
| Basidiomycota | Agaricomycetes | Russulales | Russulaceae | Russula | *Russula grisea* | contact-short | A | J | |
| Basidiomycota | Agaricomycetes | Russulales | Russulaceae | Russula | *Russula ionochlora* | contact-short | J, A | J, A | J, A |
| Basidiomycota | Agaricomycetes | Russulales | Russulaceae | Russula | *Russula nobilis* | contact-short | J, A | J, A | J, A |
| Basidiomycota | Agaricomycetes | Russulales | Russulaceae | Russula | *Russula ochroleuca* | contact-short | J, A | J, A | J, A |
| Basidiomycota | Agaricomycetes | Russulales | Russulaceae | Russula | *Russula parazurea* | contact-short | J, A | A | J, A |
| Basidiomycota | Agaricomycetes | Russulales | Russulaceae | Russula | *Russula peckii* | contact-short | J | | |
| Basidiomycota | Agaricomycetes | Russulales | Russulaceae | Russula | *Russula puellaris* | contact-short | J, A | A | |
| Basidiomycota | Agaricomycetes | Russulales | Russulaceae | Russula | *Russula recondita* | contact-short | | J | |
| Basidiomycota | Agaricomycetes | Russulales | Russulaceae | Russula | *Russula risigallina* | contact-short | | J, A | |
| Basidiomycota | Agaricomycetes | Russulales | Russulaceae | Russula | *Russula variata* | contact-short | | J, A | |
| Basidiomycota | Agaricomycetes | Russulales | Russulaceae | Russula | *Russula violeipes* | contact-short | J | | |
| Basidiomycota | Agaricomycetes | Russulales | Russulaceae | Russula | *Russula virescens* | contact-short | J | | |
| Basidiomycota | Agaricomycetes | Russulales | Russulaceae | Russula | | contact-short | J, A | J, A | J, A |
| Basidiomycota | Agaricomycetes | Russulales | Russulaceae | Russula | | contact-short | | J | |
| Basidiomycota | Agaricomycetes | Russulales | Russulaceae | Russula | | contact-short | J, A | | |
| Basidiomycota | Agaricomycetes | Russulales | Russulaceae | Russula | | contact-short | A | | |
| Basidiomycota | Agaricomycetes | Russulales | Russulaceae | | | contact-short | | J, A | J, A |
| Basidiomycota | Agaricomycetes | Sebacinales | Sebacinaceae | Sebacina | *Sebacina epigaea* | contact-short | | J, A | |
| Basidiomycota | Agaricomycetes | Sebacinales | Sebacinaceae | Sebacina | *Sebacina incrustans* | contact-short | J, A | A | J, A |

| | | | | | | | | | |
|---|---|---|---|---|---|---|---|---|---|
| Basidiomycota | Agaricomycetes | Sebacinales | Sebacinaceae | Sebacina | unidentified | contact-short | J, A | J, A | J, A |
| Basidiomycota | Agaricomycetes | Thelephorales | Thelephoraceae | Pseudotomentella | *Pseudotomentella tristis* | medium-long | A | J, A | A |
| Basidiomycota | Agaricomycetes | Thelephorales | Thelephoraceae | Pseudotomentella | | medium-long | | A | |
| Basidiomycota | Agaricomycetes | Thelephorales | Thelephoraceae | Thelephora | unidentified | medium-long | J | J, A | J, A |
| Basidiomycota | Agaricomycetes | Thelephorales | Thelephoraceae | Tomentella | *Tomentella atroarenicolor* | medium-long | J | J, A | J, A |
| Basidiomycota | Agaricomycetes | Thelephorales | Thelephoraceae | Tomentella | *Tomentella badia* | medium-long | J, A | J, A | |
| Basidiomycota | Agaricomycetes | Thelephorales | Thelephoraceae | Tomentella | *Tomentella botryoides* | medium-long | J, A | | |
| Basidiomycota | Agaricomycetes | Thelephorales | Thelephoraceae | Tomentella | *Tomentella bryophila* | medium-long | | J, A | A |
| Basidiomycota | Agaricomycetes | Thelephorales | Thelephoraceae | Tomentella | *Tomentella coerulea* | medium-long | | J, A | J, A |
| Basidiomycota | Agaricomycetes | Thelephorales | Thelephoraceae | Tomentella | *Tomentella galzinii* | medium-long | | J, A | J, A |
| Basidiomycota | Agaricomycetes | Thelephorales | Thelephoraceae | Tomentella | *Tomentella lapida* | medium-long | A | A | |
| Basidiomycota | Agaricomycetes | Thelephorales | Thelephoraceae | Tomentella | *Tomentella lilacinogrisea* | medium-long | | J, A | A |
| Basidiomycota | Agaricomycetes | Thelephorales | Thelephoraceae | Tomentella | *Tomentella papuae* | medium-long | J, A | J, A | J, A |
| Basidiomycota | Agaricomycetes | Thelephorales | Thelephoraceae | Tomentella | *Tomentella pyrolae* | medium-long | | A | |
| Basidiomycota | Agaricomycetes | Thelephorales | Thelephoraceae | Tomentella | *Tomentella stuposa* | medium-long | J | J, A | J, A |
| Basidiomycota | Agaricomycetes | Thelephorales | Thelephoraceae | Tomentella | *Tomentella subclavigera* | medium-long | J, A | J, A | J |
| Basidiomycota | Agaricomycetes | Thelephorales | Thelephoraceae | Tomentella | *Tomentella terrestris* | medium-long | J, A | J, A | J, A |
| Basidiomycota | Agaricomycetes | Thelephorales | Thelephoraceae | Tomentella | unidentified | medium-long | | A | |
| Basidiomycota | Agaricomycetes | Thelephorales | Thelephoraceae | Tomentella | | medium-long | J | J, A | J, A |
| Basidiomycota | Agaricomycetes | Thelephorales | Thelephoraceae | Tomentella | | medium-long | J, A | J, A | J |
| Basidiomycota | Agaricomycetes | Thelephorales | Thelephoraceae | Tomentellopsis | *Tomentellopsis echinospora* | medium-long | | J, A | |
| Basidiomycota | Agaricomycetes | Thelephorales | Thelephoraceae | | | medium-long | J, A | J, A | J, A |
| Ascomycota | Dothideomycetes | Mytilinidales | Gloniaceae | Cenococcum | unidentified | contact-short | J, A | J, A | J, A |
| Ascomycota | Dothideomycetes | Mytilinidales | Gloniaceae | Cenococcum | | contact-short | A | J | A |
| Ascomycota | Dothideomycetes | Pleosporales | Leptosphaeriaceae | Acicuseptoria | *Acicuseptoria rumicis* | unknown | | | J, A |
| Ascomycota | Eurotiomycetes | Eurotiales | Elaphomycetaceae | Elaphomyces | *Elaphomyces decipiens* | contact-short | J, A | | |
| Ascomycota | Eurotiomycetes | Eurotiales | Elaphomycetaceae | Elaphomyces | *Elaphomyces granulatus* | contact-short | J | | |
| Ascomycota | Eurotiomycetes | Eurotiales | Elaphomycetaceae | Elaphomyces | *Elaphomyces leucosporus* | contact-short | J | | |
| Ascomycota | Eurotiomycetes | Eurotiales | Elaphomycetaceae | Elaphomyces | *Elaphomyces muricatus* | contact-short | J, A | J, A | J, A |
| Ascomycota | Eurotiomycetes | Eurotiales | Elaphomycetaceae | Elaphomyces | *Elaphomyces papillatus* | contact-short | J | | |
| Ascomycota | Eurotiomycetes | Eurotiales | Elaphomycetaceae | unidentified | unidentified | contact-short | J, A | | J, A |
| Ascomycota | Leotiomycetes | Helotiales | Leotiaceae | Leotia | *Leotia lubrica* | contact-short | J, A | J, A | A |
| Ascomycota | Pezizomycetes | Pezizales | Discinaceae | Hydnotrya | *Hydnotrya tulasnei* | medium-long | J, A | | |
| Ascomycota | Pezizomycetes | Pezizales | Helvellaceae | Helvella | *Helvella lacunosa* | contact-short | | J, A | |
| Ascomycota | Pezizomycetes | Pezizales | Pezizaceae | Hydnobolites | *Hydnobolites cerebriformis* | contact-short | J | A | J, A |
| Ascomycota | Pezizomycetes | Pezizales | Pezizaceae | Pachyphloeus | unidentified | medium-long | | J, A | J |
| Ascomycota | Pezizomycetes | Pezizales | Pyronemataceae | Genea | *Genea darii* | contact-short | | J, A | |
| Ascomycota | Pezizomycetes | Pezizales | Pyronemataceae | Genea | *Genea hispidula* | contact-short | | J, A | |

| | | | | | | | | | |
|---|---|---|---|---|---|---|---|---|---|
| Ascomycota | Pezizomycetes | Pezizales | Pyronemataceae | Genea | unidentified | contact-short | | J, A | |
| Ascomycota | Pezizomycetes | Pezizales | Pyronemataceae | Otidea | *Otidea bufonia* | unknown | | A | |
| Ascomycota | Pezizomycetes | Pezizales | Pyronemataceae | Otidea | *Otidea onotica* | unknown | | J, A | |
| Ascomycota | Pezizomycetes | Pezizales | Pyronemataceae | Tarzetta | *Tarzetta catinus* | unknown | | J, A | A |
| Ascomycota | Pezizomycetes | Pezizales | Tuberaceae | Tuber | *Tuber borchii* | contact-short | J | J, A | A |
| Ascomycota | Pezizomycetes | Pezizales | Tuberaceae | Tuber | *Tuber puberulum* | contact-short | | J, A | J |
| Ascomycota | Endogonomycetes | Endogonales | Endogonaceae | Endogone | *Endogone lactiflua* | unknown | J | | |

**Table S3.** Results of PERMANOVA analysis to test the effect of site (Northern, Intermediate and Southern) and sampling campaign (spring, June, and summer, August) on the ectomycorrhizal fungal communities. Df: degrees of freedom; Sum Sq: sum of squares; Pseudo-*F*: *F*-value by permutation, *P*(perm): *P*-values based on 9,999 permutations. Statistically significant *P*-values are shown in bold.

| **Source** | **Df** | **Sum Sq** | **Pseudo-*F*** | $R^2$ | ***P*(perm)** |
|---|---|---|---|---|---|
| Site | 2 | 3.08 | 4.88 | 0.18 | **<0.001** |
| Campaign | 1 | 0.22 | 0.69 | 0.01 | 0.95 |
| Site × Campaign | 2 | 0.48 | 0.75 | 0.03 | 0.97 |
| Residuals | 42 | 13.28 | | 0.78 | |
| Total | 47 | 17.06 | | 1.00 | |

**Table S4.** Results of the linear mixed models to test for the effect of topsoil (0-5 cm) volumetric water content (VWC) across sampling campaigns (spring, June, and summer, August) and their interaction in: species richness and species diversity of ectomycorrhizal fungi. Marginal ($R^2_m$) and conditional ($R^2_c$) coefficients indicate the variance explained by fixed only and by fixed and random factors, respectively. Df: degrees of freedom; Sum Sq: sum of squares. Statistically significant *P*-values are shown in bold.

| **Species richness** | | | | | |
|---|---|---|---|---|---|
| **Source** | **Df** | ***F*** | ***P*** | $R^2_m$ | $R^2_c$ |
| VWC | 1 | 0.77 | 0.387 | 0.09 | 0.09 |
| Campaign | 1 | 2.37 | 0.135 | | |
| VWC × Campaign | 1 | 1.28 | 0.268 | | |
| **Species diversity** | | | | | |
| **Source** | **Df** | ***F*** | ***P*** | $R^2_m$ | $R^2_c$ |
| VWC | 1 | 4.48 | **0.044** | 0.18 | 0.18 |
| Campaign | 1 | 2.07 | 0.162 | | |
| VWC × Campaign | 1 | 0.38 | 0.542 | | |

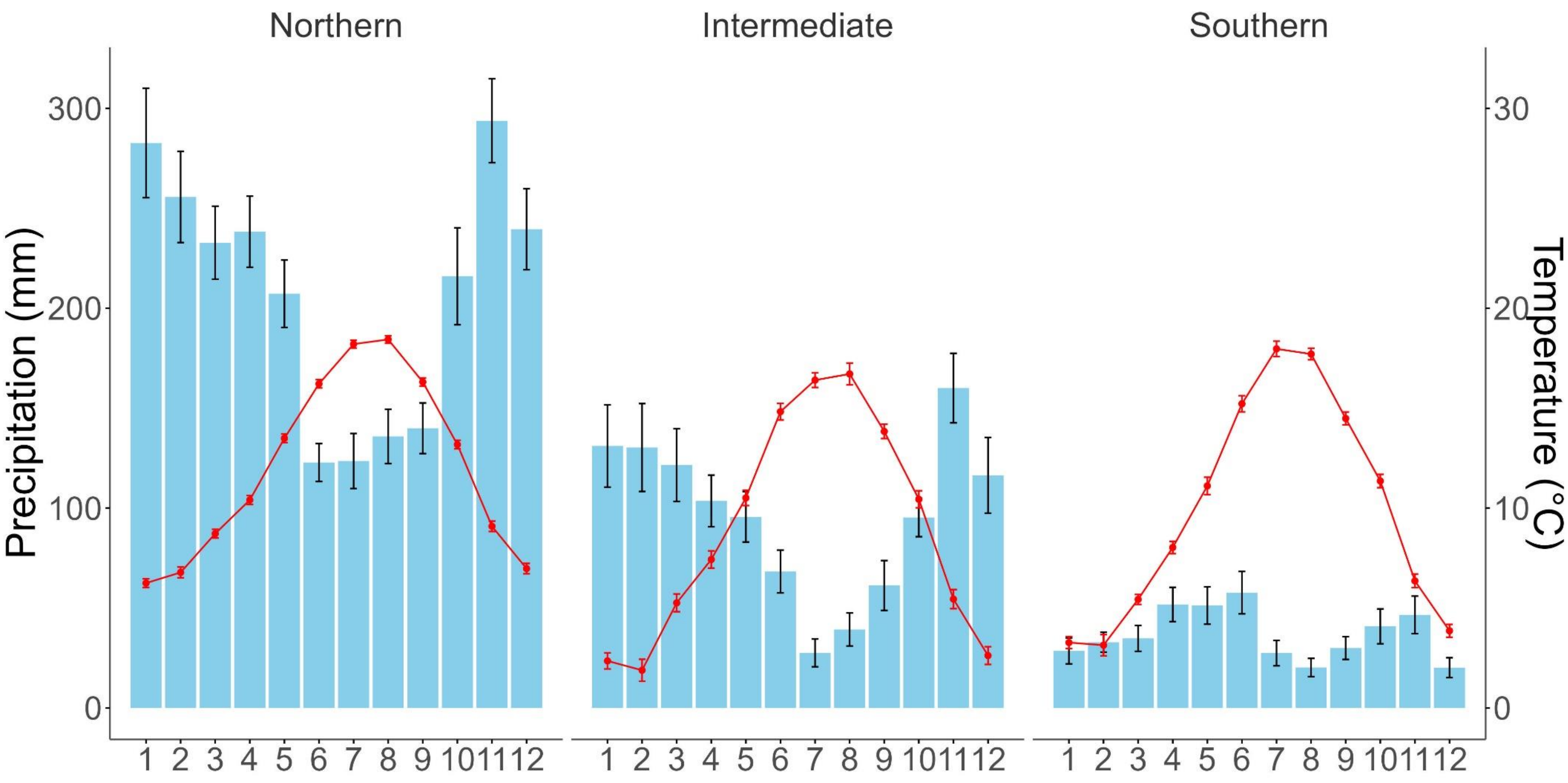


**Figure S1.** Mean (± se, *n* varies depending on the length of the time series, see methods) monthly precipitation (blue bars) and temperature (red lines) for each of the three study sites in Northern Spain: Northern (Artikutza, Navarra), Intermediate (Iturrieta, Álava) and Southern (Diustes, Soria).

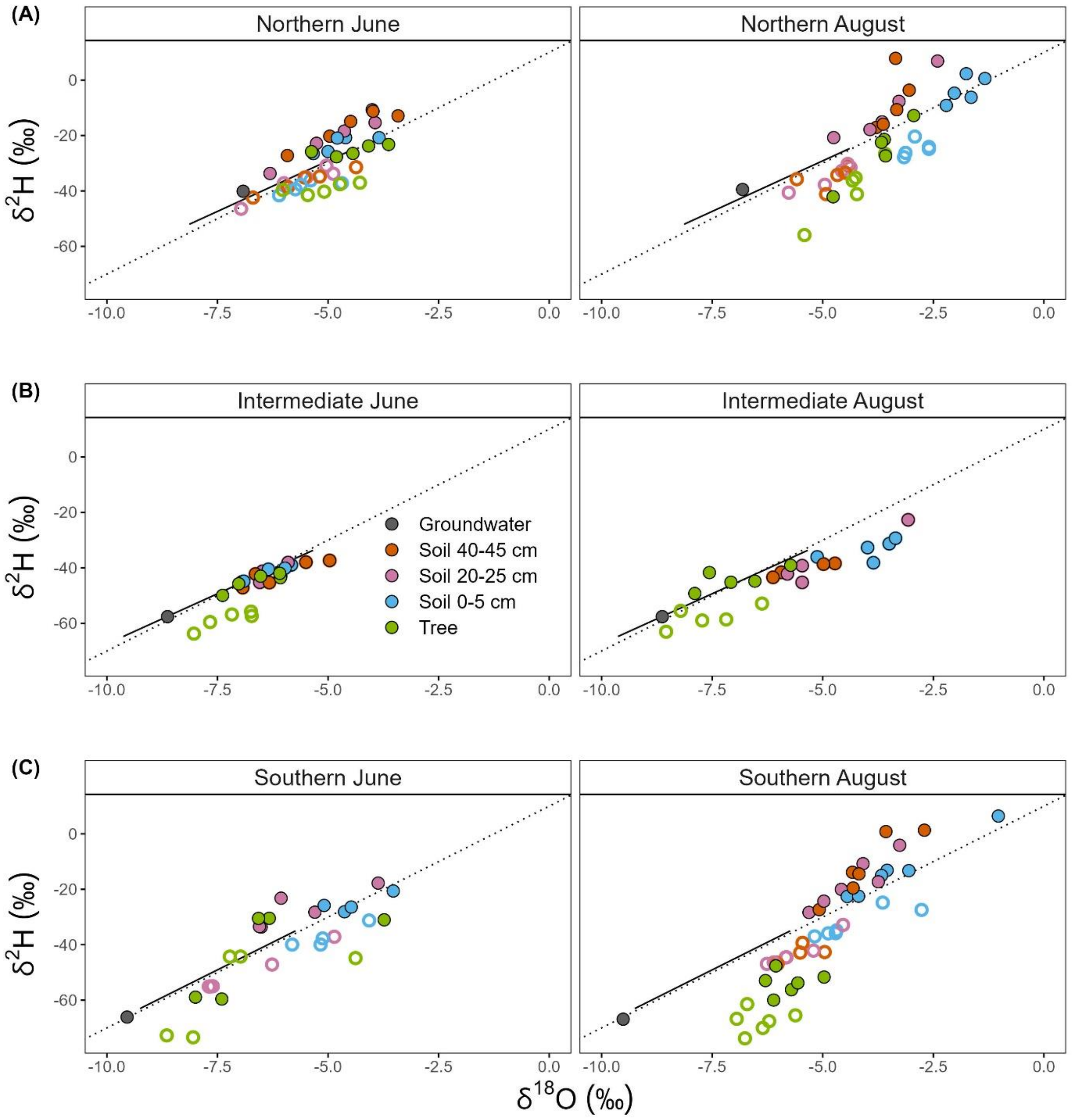


**Figure S2.** Dual-isotope ($\delta^2H$ vs. $\delta^{18}O$) plots of tree water (green symbols), soil water (blue, purple and orange symbols) and groundwater (GW, grey symbols) for the three study sites (A-C) and both sampling campaigns (spring, June, and summer, August). Closed green circles correspond to estimated tree sap water and open green circles to bulk tree water. Closed blue, purple and orange circles correspond to expected plant available water: unbound soil water at the clayey sites (Northern and Southern), and bulk soil water at the sandy site (Intermediate); open blue, purple and orange circles at the Northern and Southern sites represent isotopic composition of bulk soil water. Lines are the global meteoric water line (dotted line, $\delta^2H = 10 + 8\ \delta^{18}O$) and the local meteoric water lines (solid lines, slope and intercept specific for each site obtained from Nelson et al. 2021).

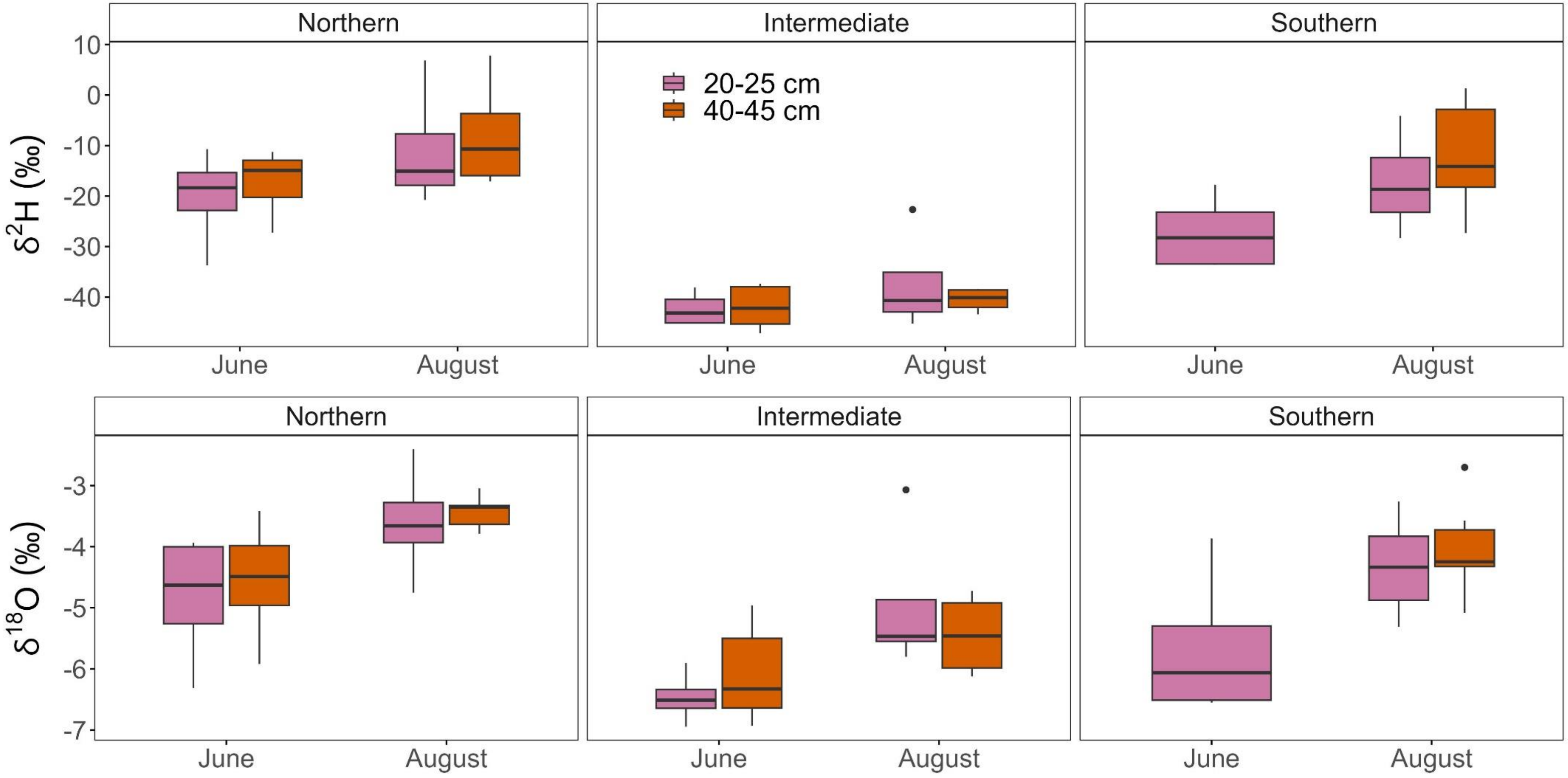


**Figure S3.** Boxplots of hydrogen ($\delta^2$H) and oxygen isotopic composition ($\delta^{18}$O) of 20-25 cm and 40-50 cm soil water, for the three study sites and two sampling campaigns (spring, June, and summer, August). No 40-45 cm soil samples were collected at the Southern site in the June campaign. Depicted soil water isotopic compositions are those estimated for unbound soil water (for the Northern and Southern sites) or bulk soil water (for the Intermediate site).
